# RADIP technology comprehensively identifies H3K27me3-mediated RNA-chromatin interactions

**DOI:** 10.1101/2024.06.04.597497

**Authors:** Xufeng Shu, Masaki Kato, Satoshi Takizawa, Yutaka Suzuki, Piero Carninci

## Abstract

Many RNAs associate with chromatin, either directly or indirectly. Several technologies for mapping regions where RNAs interact across the genome have been developed to investigate the function of these RNAs. Obtaining information on the proteins involved in these RNA–chromatin interactions is critical for further analysis. Here, we developed RADIP (RNA and DNA interacting complexes ligated and sequenced (RADICL-seq) with immunoprecipitation), a novel technology that combines RADICL-seq technology with chromatin immunoprecipitation to characterize RNA–chromatin interactions mediated by individual proteins. Building upon the foundational principles of RADICL-seq, RADIP extends its advantages by increasing genomic coverage and unique mapping rate efficiency compared to existing methods. To demonstrate its effectiveness, we applied an anti-H3K27me3 antibody to the RADIP technology and generated libraries from mouse embryonic stem cells (mESCs). We identified a multitude of RNAs, including RNAs from protein-coding genes and non-coding RNAs, that are associated with chromatin via H3K27me3 and that likely facilitate the spread of Polycomb repressive complexes over broad regions of the mammalian genome, thereby affecting gene expression, chromatin structures and pluripotency of mESCs. Our study demonstrates the applicability of RADIP to investigations of the functions of chromatin-associated RNAs.

**GRAPHICAL ABSTRACT:** 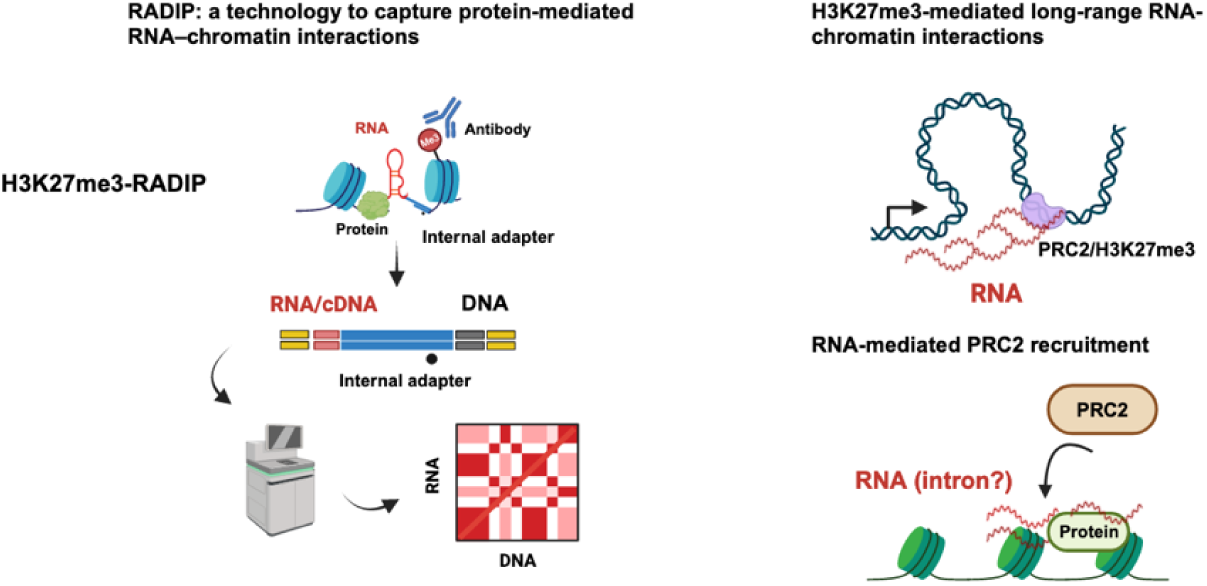

## INTRODUCTION

Large portions of mammalian genomes are transcribed into RNAs (1–3). Many of these RNAs are known to interact with chromatin, either directly or indirectly, and they are typically identified as chromatin-associated RNAs (caRNAs). caRNAs include many long non-coding RNAs (lncRNAs) and play essential roles in gene regulation, genomic organization, and chromatin structure maintenance (4–6).

Several technologies have been developed to determine the functional roles of caRNAs and lncRNAs. RNA-centric methods (e.g., CHART, ChiRP, and RAP) (7–9) for studying RNA–chromatin interactions have been used for a while. However, the reliance of these technologies on antisense probes to target specific transcripts limits their capacity to capture RNA–chromatin interactions within the nucleus comprehensively. More recently, some technologies that are based on a proximity ligation method of mapping genome-wide RNA–chromatin interactions have been developed; they include GRID-seq, iMARGI, ChAR-seq, RADICL-seq, and Red-C (10–14). In addition, SPRITE and its enhanced successor, RD-SPRITE (15,16), which function by breaking crosslinked nuclei into multiple particles by sonication and tagging the identical unique barcodes to all RNA and DNA molecules within the same subnuclear particle, have been used to investigate RNA–DNA interactions. However, very deep sequencing is required to fully identify all RNA–chromatin interactions owing to a wide range of RNA expression levels. Although these methods can broadly detect the RNA–chromatin interactome, they can not broadly reveal which proteins or chromatin modifications are associated with specific RNAs. Therefore, understanding information on the proteins involved in RNA-chromatin interactions is essential for deepening the analysis.

Polycomb repressive complex (PRC) 2 is a histone-modifying complex responsible for the trimethylation of histone H3 lysine 27 marks (H3K27me3), associated with the repression of cell-type-specific genes. It has been postulated that many lncRNAs interact with PRC2 to facilitate the deposition of H3K27me3 marks and thus regulate the expression of target genes. For instance, *XIST* (X-inactive-specific transcripts) and *HOTAIR* (HOX transcript antisense intergenic RNA) lncRNAs can bind to PRC2 and then recruit and guide the PRC2 to specific genomic regions for target gene silencing (17,18).

However, our understanding of the mechanisms of PRC2 regulation by RNA is controversial. Recent studies have shown that PRC2 interacts with RNAs, including RNAs derived from bacterial sequences, with little specificity, in a manner described as “promiscuous” (19,20). These spurious interactions raise the question of whether lncRNAs can provide specificity for PRC2 localization on the genome. Other studies have reported that RNAs can inhibit PRC2 catalytic activity by competing with chromatin for PRC2 binding (21–26). In contrast, another study demonstrated that PRC2–RNA interaction is critical *in vivo* for PRC2 chromatin occupancy and localization across the genome and for maintaining H3K27me3 levels in induced pluripotent stem cells (27).

Many lncRNAs engage in long-range RNA–chromatin interactions (9, 17, 28). Several studies have revealed that PRCs establish long-range DNA–DNA interactions in various types of cells (29–33). However, the precise mechanisms of how RNAs influence the establishment and organization of the three-dimensional genome architecture mediated by PRCs or H3K27me3 histone marks remain unclear.

The study of caRNAs and lncRNAs linked to PRCs and H3K27me3 histone marks using the above methods is hampered by the scarcity of these RNAs, and their function may be repressive. To enable the characterization of the proteins that are involved in RNA–chromatin interactions, we have developed RADIP (RNA and DNA interacting complexes ligated and sequenced (RADICL-seq) with immunoprecipitation), a novel technology that inherits advantages of RADICL-seq methods by broadly determining RNA–chromatin interactions, in conjunction with chromatin immunoprecipitation (ChIP), to identify those interactions mediated by a specific protein or histone mark of interest. We used an anti-H3K27me3 antibody to study the H3K27me3-mediated RNA–chromatin interactome in mouse embryonic stem cells (mESCs), and we demonstrated the applicability of the RADIP method to investigations of the function of caRNAs. We identified a spectrum of caRNAs enriched at, and around, PRC2-binding chromatin sites and H3K27me3 peak regions in mESCs; these caRNAs may be involved in PRC2-mediated gene repression and the establishment of PRC2-mediated three-dimensional genome architecture in mESCs.

## MATERIALS AND METHODS

### Cell culture and crosslinking of cells

mESCs (PGK12.1) (34) were cultured in Dulbecco’s modified Eagle’s medium (Nacalai Tesque) supplemented with 15% fetal bovine serum (Gibco), 0.1 mM 2-mercaptoethanol (Gibco), 1,000 U/mL Leukemia Inhibitory Factor (Merck), and 1×□non-essential amino acids (Gibco). Detached cells were pelleted and resuspended in fresh 1% formaldehyde (Thermo Fisher Scientific) at a volume of 1 mL formaldehyde for 2 million cells. Cells were incubated at room temperature for 10 min with rotation. Glycine (Sigma–Aldrich) was added at a final concentration of 125 mM to quench the formaldehyde, and the cells were incubated at room temperature for 5 min with rotation. Cells were pelleted and washed with ice-cold phosphate-buffered saline (PBS), pelleted again, snap frozen in liquid nitrogen and stored at –80 □.

### RADIP library preparation

#### Generation of chimeric molecules of RNA – internal bridge adaptor – DNA

Chimeric molecules of RNA – internal bridge adaptor –DNA were generated following a published protocol, with modifications (12), except without using AMPure XP magnetic beads. In brief, 20 million crosslinked cells were resuspended in 4 mL of ice-cold lysis buffer (10 mM Tris-HCl pH 8.0, 10 mM NaCl, 0.2% Igepal CA-630 (octylphenoxy poly(ethyleneoxy)ethanol), 1 mM PMSF (phenylmethylsulfonyl fluoride), 1× cOmplete Protease Inhibitor Cocktail (Roche), 0.8 U/μL RNasin Plus (Promega)) and incubated on ice for 10 min. Chimeric molecules of RNA – internal bridge adapter –DNA were generated with 2 million cells per tube; multiple tubes were processed in parallel before the sonication step and then pooled. 0.122 U/μL RNase H (New England Biolabs) was added and the reaction was incubated at 37 °C for 40 min before the RNA – internal bridge adapter ligation step following a published protocol (12).

#### Sonication

The nuclei were pelleted at 2500□×*g* for 4 min, resuspended in 100□μL SDS (sodium dodecyl sulfate) lysis buffer (50□mM Tris-HCl pH 8.0, 1% SDS, 10 mM EDTA), and incubated on ice for 10 min. After the addition of 400 μL of cold ChIP dilution buffer (50□mM Tris-HCl pH 8.0, 167□mM NaCl, 1.1% Triton X-100, 0.1% sodium deoxycholate), the mixture was briefly sonicated (5 s at 4 □) in a Picoruptor sonicator (Diagenode). The sonicated mixture was spun at 14,000 rpm for 5 min at 4 □, and the supernatant was collected. After adding 600□μL of ChIP dilution buffer, 100 μL of the mixture was used as a 10% input sample (for the Input library), and the remainder was used for immunoprecipitation.

#### Chromatin immunoprecipitation

A rabbit monoclonal antibody against H3K27me3 (Cell Signaling Technology, #9733) was used. The antibody was incubated with anti-rabbit IgG Dynabeads (Thermo Fisher Scientific) on ice for 1 h and then overnight after the addition of the sonicated chimeric molecules at 4 □. The sample was washed with RIPA buffer (50□mM Tris-HCl pH 8.0, 500□mM NaCl, 1% Triton X-100, 0.1% NaDOC (sodium deoxycholate), 1 mM EDTA, 0.1% SDS), the original RADICL-seq protocol (12) was followed for the rest of the procedure (for the H3K27me3-RADIP library).

### Data processing

#### RADIP mapping and processing

RADIP mapping and processing were performed as described before (12). A summary of the pipeline workflow is depicted in Supplementary Figure S1A.

#### Reference annotation and genome binning

The comprehensive gene annotation for mouse (GENCODE release M25 version for mm10 assembly) (35) was downloaded. Genome binning was performed as described before (12). Mouse genome binning at 25-kb resolution was performed using BEDtools (ver. 2.26.0) (36) with parameter -w 25000. RNA and DNA tags were associated unambiguously with the corresponding annotated features (RNA tags were mapped to reference annotated genes, and DNA tags were mapped to genomic 25-kb bins). We identified each fragment by the single-nucleotide position at its center to reduce the possibility of fragments overlapping with multiple genes or bins. The strand information was considered for RNA tags but discarded for DNA tags. Both RNA and DNA tags were required to be mapped uniquely [Burrows–Wheeler Aligner MAPping Quality (BWA MAPQ)□=□37] to the genome.

#### Reproducibility

The reproducibility of the RNA–DNA interaction frequencies across replicates was assessed by counting the occurrences of genic transcripts and 25-kb bin interaction pairs. The results were visualized using the CAGEr (ver. 2.4.0) R package (37).

#### Removal of interaction pairs from blacklisted genomic regions

A blacklist of the mouse genome (mm10) was downloaded from the ENCODE Data Coordination Center. All RNA–DNA interaction pairs associated or overlapped, or both, with these blacklist mouse mm10 genomic regions were removed by using the BEDtoolsr (ver. 2.30.0-5) R package (38) based on the genomic coordinates of the DNA tags.

#### ChIP-seq and ATAC-seq data processing

EZH2 and H3K27me3 ChIP-seq data were obtained from the GEO (Gene Expression Omnibus) database (GSM5658769 and GSM5658773, respectively). H3K9me3 ChIP-seq and ATAC-seq data were obtained from the GEO database (GSM5233300 and GSM5582921, respectively). DNA tags from RADIP RNA–DNA interactions were extracted, and then the distributions of the DNA tags around the center points of the H3K27me3 ChIP peaks, the center points of the EZH2-binding sites, the center points of the H3K9me3 ChIP peaks and the center points of the ATAC peaks were computed by using the computeMatrix function of deepTools (ver. 3.5.0) (39).

#### High-confidence interaction calculation and definition or background removal step

A modified negative binomial distribution approach based on the CHiCANE R package (ver. 0.1.8) (40) was used to calculate the adjusted *P* value (*q* value) for each RNA–DNA interaction pair. Only statistically significant interaction pairs (*q* value ≤ 0.1) were retained for subsequent analysis.

#### Genome-wide RNA–DNA interaction plots

For the genome-wide interaction plots, the genome was divided into 25-kb bins, and each interaction was assigned to these bins based on both RNA and DNA tags in the significant interaction pairs. Thus, the RNA–DNA interaction pairs were changed to bin–bin interactions across the mouse mm10 genome. The plot was produced in R using the ggplot2 (ver. 3.4.2) R package (41) to visualize all interactions.

#### PAR-CLIP (EZH2 targeting) and RIP-seq (EZH2 targeting) data processing

PAR-CLIP data were obtained from the GEO database (GSE49433). The mouse mm9 genome used in the PAR-CLIP data was liftOvered to the mouse mm10 genome using CrossMap (ver. 0.5.2) (42). RIP-seq data were obtained from the GEO database (GSM4080157). To enable us to access the genomic coordinates more efficiently, the BigWig format was changed to a BED-based format by bigWigtoBedGraph, a tool released on the UCSC (University of California Santa Cruz) genome browser. The processed PAR-CLIP and RIP-seq data were merged to define EZH2- or PRC2-interacting RNAs.

#### Differential interaction analysis

Common RNAs were determined from RNA names, and RNA–DNA interactions were counted per RNA name and TMM normalized by using edgeR (ver. 3.40.2) (43). A volcano plot was created based on the results of the differential interaction analysis after using edgeR (ver. 3.40.2) with customized cutoffs (adjusted *P* ≤ 0.1 and absolute value of log2 fold change ≥ 0.5).

#### RNA-seq data processing

Enriched RNAs’ expression values (in RPKM, reads per kilobase per million mapped reads) were retrieved from the mESCs RNA-seq data (GEO: GSM3239395). EZH2 knockout and EZH2 wild-type RNA-seq reads were obtained from GEO (GSE66814) and were mapped to the mm10 genome by using STAR (ver. 2.5.0a) (44). They were then intersected with GENCODE ver. M25 to generate counts by using htseq-count (ver. 0.13.5) (45). All counts were normalized using the DESeq2 (ver. 1.38.3) R package (46) with the default settings. The log2 fold-change values of the normalized RNA-seq data were calculated. Significance was accessed using the ggsignif (ver. 0.6.4) R package (47).

#### GO term analysis

The shift from RNA names to RNA Entrez IDs was done using *bitr function* in clusterProfiler (ver. 4.6.2) (48) R package. The enrichGO function was used to call biological processes significantly associated with the input query RNAs, using the default settings.

#### TLDs

TLDs were built on the basis of *cis* RNA–DNA interaction pairs on each chromosome from chr. 1 to chr. 19 and chr. X by using SuperTAD (ver. 1.2) with the recommend settings (49). After an overall comparison, the TLDs that had changed across the whole mouse mm10 genome were defined and the RNAs located in both anchors of the changed TLDs in the H3K27me3-RADIP samples were defined as “Changed TLD” RNAs.

#### Hi-C paired-end-read processing and comparison with RADIP RNA-DNA interactions

Publicly available Hi-C data for mESCs were obtained (GEO: GSE137274). The Hi-C data were analyzed by using Juicer (ver. 1.6) and Juicer Tools (ver. 1.9.9) (50) with MAPQ□=□30. The processed Hi-C data were binned at a fixed resolution of 25 kb and KR (Knight–Ruiz) normalized by using straw in Juicer Tools, which is a data extraction API (application program interface) for Hi-C contact maps. Pearson’s correlation coefficient was calculated by comparing Hi-C DNA–DNA contacts with RNA–DNA interactions at the same 25-kb resolution. The TAD structures were also called by using Juicer Tools at 25-kb resolution. Visualization was done by using CoolBox (ver. 0.3.8) (51) and ggplot2 (ver. 3.4.2).

#### Elucidation of the RNA-induced genomic distribution of H3K27me3 and its impact on gene expression

The publicly available data of a differentiation model was utilized. This model involved the culture of mESCs under conditions with leukemia inhibitory factor (LIF) and induced differentiation into embryoid bodies following the removal of LIF and supplementation with 10 µM retinoic acid (RA) for 6 days. RNA-seq and ChIP-seq data were downloaded at the day 0 and 6 time points (GEO: GSE169745). RNA-seq and ChIP-seq data were processed as described above.

#### Sequence motif enrichment analysis

The STREME package (ver. 5.5.3) (52) was used to identify motifs enriched in RNA tags in RADIP libraries. RNA tags of interest were collected and extended by 800 bp from the central points in either direction to simulate the average length of the ChIP peaks for better STREME performance and to overcome the fact that our RNA tags were only about 27 nt long. Ungapped motifs enriched in 1.6-kb query sequences were discovered by using STREME with custom cutoffs (*P* value was no more than 0.05 and enriched peaks were near the central point, standing for the real captured RNA tags containing the identified motifs). The FIMO package (ver. 5.5.3) (53) was used to find those submitted RNA sequences that contained the selected motifs.

## RESULTS

### RADIP technology workflow

We developed the RADIP method to identify RNA–chromatin interactions mediated by a particular protein or specific histone mark by combining the RADICL-seq protocol with a chromatin immunoprecipitation protocol (Figure 1A, see Materials and methods). The nuclei of mammalian cells crosslinked in 1% formaldehyde were isolated, and their genomic DNA was digested with DNase I. We followed the RADICL-seq protocol (12) until the step of RNA–chromatin chimera formation, except omitting the use of AMPure XP magnetic beads due to the difficulty in solubilizing RNA–chromatin chimeras once they have attached to the beads. The nuclear membrane was disrupted by sonication, but the sonication time was kept brief to avoid damaging the RNA. Immunoprecipitation was performed using an antibody against a particular protein or a specific type of histone mark with RNA–adaptor–DNA chimeric molecules. The antibody had selectively captured the target chimera to prepare the RADIP samples, and excess uncaptured molecules were washed away. In parallel, we kept ten percent of RNA–adaptor–DNA chimeric molecules and prepared libraries without antibody selection as the Input sample for comparison. The Input sample essentially corresponds to the original RADICL-seq method (therefore by omitting antibody selection), which we use to calculate the enrichment or depletion of specific RNAs in the RADIP interactomes. For convenience, in subsequent analysis, we refer to it simply as the “Input” sample. To comprehensively investigate the functions of RNAs interacting with genomic sites marked by H3K27me3, we utilized H3K27me3-RADIP to examine the H3K27me3-mediated RNA–chromatin interactome in mouse embryonic stem cells (mESCs). We demonstrate the importance of RADIP-seq by elucidating which RNAs collaborate with PRC2 to regulate specific gene expressions and maintain chromatin structures.

**Figure 1.**
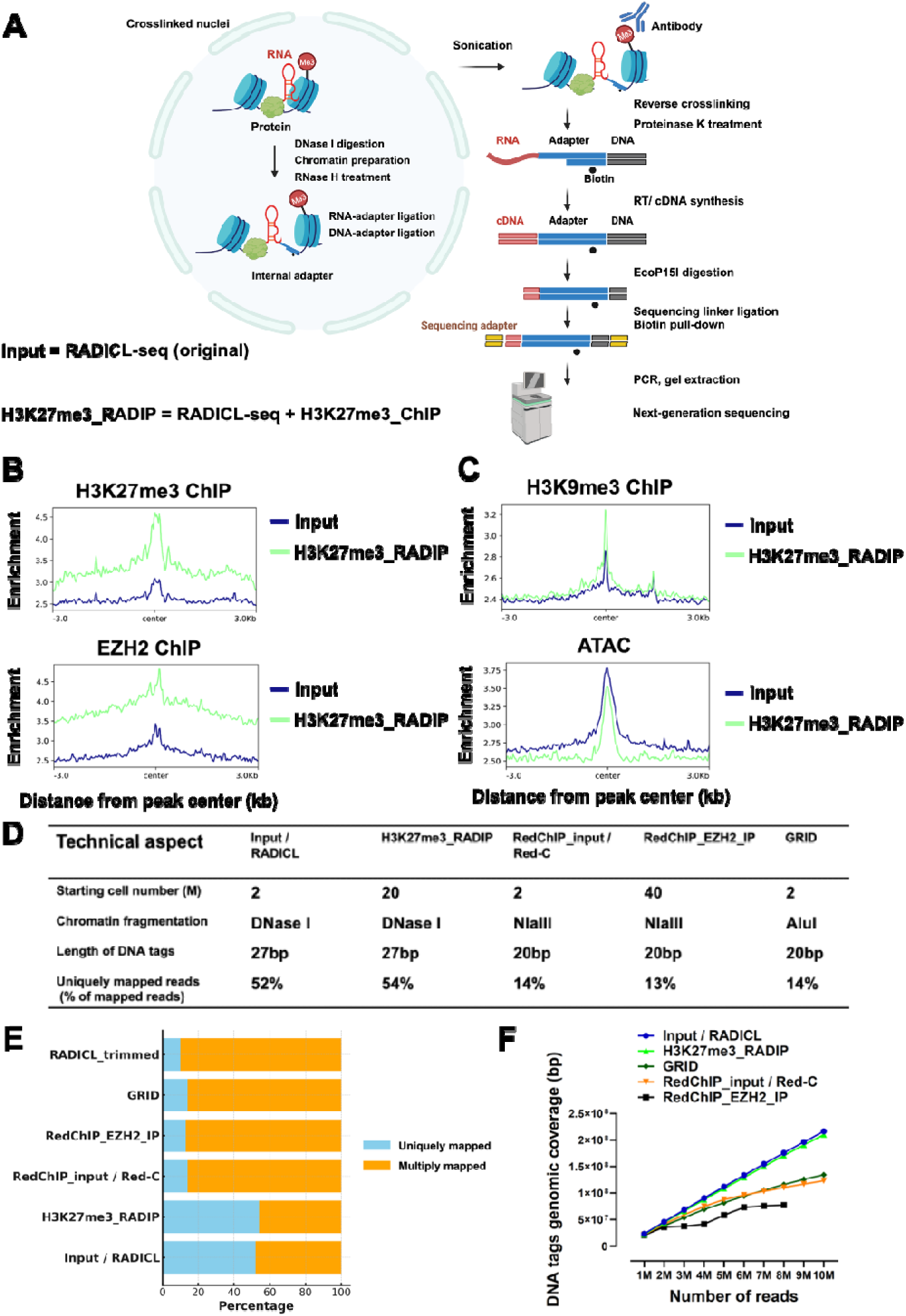
RADIP technology enables the capture of RNA–chromatin interactions mediated by a specific protein or certain types of histone marks. (A) Schematic of the RADIP technology. Left: steps performed i*n situ* on fixed nuclei after partial lysis of the nuclear membrane. Right: steps performed in solution after disruption of the nuclear membrane by sonication. After reversal of crosslinks, the RNA–adaptor–DNA chimera is converted into a fully double-stranded DNA (dsDNA) molecule. This molecule is then digested to a designated length using the EcoP15I enzyme. (B) Distribution of DNA tags from Input and H3K27me3-RADIP samples around H3K27me3 and EZH2 ChIP-seq peaks in local windows (± 3 kb from the center points of the ChIP-seq peaks). (C) Distribution of DNA tags from Input and H3K27me3-RADIP samples around H3K9me3 ChIP-seq peaks and ATAC-seq peaks in local windows (± 3 kb from the center points of the ChIP-seq peaks and ATAC-seq peaks). (D) Comparison of features across different technologies. (E) Analysis of read length and mapping results includes uniquely mapped (blue) and multiply mapped (orange) reads reported as percentages of the total read pool. For direct comparison, both DNA and RNA tags of RADICL-seq reads were artificially trimmed to 20 nucleotides. (F) Assessment of genomic coverage of DNA tags across different technologies in relation to sequencing depth. All panels were generated by using datasets before the background removal step.

### Quality assessment of RADIP library

We prepared libraries of four biological replicates from the Input samples and four from the H3K27me3-RADIP samples. After rRNA removal, genome alignment, and filtering, uniquely mapped RNA–DNA pair reads were used (Supplementary Figure S1A). RADIP exhibited high reproducibility across replicates from each set of samples (Supplementary Figure S1B). We, therefore, merged all replicates in each condition to obtain 270 million 150-nt single-end raw reads from the Input library and 175 million 150-nt single-end raw reads from the H3K27me3-RADIP library. First, we examined the length and sequencing quality of RNA tags and DNA tags 25–27 nt long since the EcoP15I enzyme was used to obtain a designated length of the library (Supplementary Figure S2A and S2B). We obtained high sequencing quality, with more than 99% base-call accuracy (Phred score ≥ 25; Supplementary Figure S2C). The distribution of nucleotides in the RNA and DNA tags was uniform, except for the overhang of the T and EcoP15I recognition motif on the two ends of the DNA tags (Supplementary Figure S2D).

After the initial mapping, we used uniquely mapped RNA–DNA pair reads to assess the immunoprecipitation efficiency. Unlike the DNA tags from the Input reads, the DNA tags from the H3K27me3-RADIP reads were preferentially localized at the H3K27me3 ChIP-seq peak regions and at the binding sites of an enzyme that participates in histone methylation and ultimately in transcriptional repression, namely Enhancer of zeste homolog 2 (EZH2) (Figure 1B), indicating that the H3K27me3-RADIP technology preferentially enriched specific fractions within the initial RADICL-seq-derived data. This demonstrates the successful enrichment for PRC2-associated RNA–DNA interactions. In addition, we observed that DNA tags from the H3K27me3-RADIP reads were more enriched in heterochromatic regions marked by H3K9me3, and less enriched in open chromatin regions, as indicated by ATAC-seq data, compared to the DNA tags from the Input reads (Figure 1C).

### Comparison of RADIP with existing technologies

Building upon the foundational principles of RADICL-seq, RADIP extends its advantages by enhancing genomic coverage and increasing the efficiency of unique mapping rates compared to existing methods. These improvements are realized through two main technical aspects (Figure 1D), detailed as follows:

(i) The GRID-seq (10), Red-C (13), and RedChIP (54) protocols utilize the type II restriction enzyme MmeI to trim DNA tags from chimeric RNA–DNA interaction reads, producing a maximum of 20nt per DNA tag. In contrast, the 27nt DNA tags generated by the EcoP15I restriction enzyme used in RADICL-seq and RADIP protocols produce more extended DNA tags, which are mapped uniquely to the genome much more efficiently when compared to GRID-seq. Although the Red-C and RedChIP protocols did not trim RNA tags, the absence of size selection and purification leads to a substantial proportion of uninformative sequences in the final library. Notably, when comparing the percentage of sequencing reads that can be mapped to the mouse genome on both DNA and RNA sides, RADICL-seq and RADIP outperform GRID-seq, Red-C, and RedChIP, with more than a four-fold increase in the percentage of unique mapping reads (54% versus 13%). To confirm these results, we artificially trimmed RADICL-seq tags from 27nt to 20nt for both RNA and DNA reads and observed a dramatic reduction in the fraction of uniquely mapped RNA–DNA interactions, dropping from 52% to 10%. This reduced unique mapping rate is comparable to that observed in the GRID-seq, Red-C, and RedChIP datasets, which contain DNA tags of 20nt (Figure 1E). (ii) Unlike Red-C and RedChIP, which employ the restriction enzyme NlaIII, and GRID-seq, which uses AluI to digest genomic DNA, RADICL-seq and RADIP utilize controlled DNase I digestion. This method effectively avoids sequence biases inherent in restriction enzyme-based distribution on the genomes, achieving more uniform chromatin shearing. RADICL-seq and RADIP not only deliver higher genomic coverage, demonstrating increased coverage to sequencing depth, maintaining comparable coverage levels throughout. In contrast, starting with the same cell number and exhibiting similar unique mapping rates, GRID-seq and Red-C achieve lower coverage than RADICL-seq and RADIP, which plateaus upon analyzing 10 million RNA–DNA interactions. RedChIP, building upon Red-C technology, selectively enriches for protein-associated RNA–DNA interactions from the whole RNA–DNA interactions detected by Red-C. However, this selective enrichment further narrows the genomic regions that are finally obtained, resulting in the lowest DNA coverage among the technologies evaluated, with a coverage plateau observed at 6 million RNA–DNA interactions (Figure 1F).

These analyses suggest that RADIP comprehensively captures H3K27me3-associated or PRC2-associated RNA-chromatin interactions with reduced genomic bias compared to RedChIP. This highlights RADIP’s superior ability to thoroughly investigate protein-associated RNA–DNA interactions, thereby enhancing our understanding of the functional roles of these chromatin-bound RNAs.

### Removal of background from RNA–DNA interaction pairs in RADIP data

Previously, we showed that more than 60% of the uniquely mapped reads in the RADICL-seq data consisted of a single read interaction per bin when we spliced the linear genome into bins of 25-kb intervals (12). In our current study, we also observed many interactions in *trans* involving transcripts from protein-coding genes. Still, we could not assess whether they represented specific interactions, as speculated previously (10,13,14). Moreover, we observed many *trans* interactions in the Input data (69%) and H3K27me3-RADIP data (73%). If the RNA tags were assumed to be baits and the DNA tags were assumed to be targets, then the RADIP data generated a large volume of RNA–DNA interaction data with a complexity equivalent to that of the DNA–DNA contact data obtained by using Capture Hi-C (CHi-C) technology. We therefore adapted the CHiCANE package (40), designed for the analysis of CHi-C data, for the statistical assessment of RNA–DNA interactions (Supplementary Methods). An RNA–DNA interaction matrix based on all observed interaction pairs was assembled by individually mapping RNA tags to their reference loci and their corresponding DNA tags in 25-kb genomic bins. By considering RNA’s ability to interact with chromatin DNA, we created the background for each RNA using the truncated NB (negative binomial) test. We compared the observation of each RNA interaction with a genomic bin with the background and assigned *P* values. The false discovery rate was estimated using Benjamini–Hochberg multiple-testing correction, and the RNA–DNA interaction pairs were judged to be “significant” if the adjusted *P* value was ≤ 0.1. We removed background RNA–DNA interactions and inferred a set of 1.3 million “significant” RNA–DNA interaction pairs in the Input dataset and 1.4 million in the H3K27me3-RADIP dataset (Supplementary Figure S3A), even though many of the *trans* interactions were eliminated because of their low frequency (Supplementary Figure S3B).

### H3K27me3-RADIP enriched more long-range RNA–chromatin interactions compared to the Input

To visualize the RNA–chromatin interactions globally, we reorganized the RNA tags and their corresponding DNA tags into a two-dimensional contact matrix in 25-kb bins to represent RNA–DNA interactions in the Input and H3K27me3-RADIP samples (Figure 2A). The diagonal signal clearly showed a tendency towards proximal *cis* interactions (interactions near the genomic regions where RNAs were transcribed) in both the Input and the H3K27me3-RADIP samples. However, we observed that certain RNAs could interact with chromatin in a long-range *cis* or *trans* manner (Figure 2A). Compared with the Input samples, the H3K27me3-RADIP samples demonstrated larger numbers of long-range *cis*, which were distant from the diagonal line yet remained on the same chromosome, and increased *trans* interactions (Figure 2B). We quantified all RNA–DNA interactions according to the distance categories. We found that the interactions in the long-range *cis* (100 kb to 1 Mb and more than 1 Mb) and *trans* categories were larger in the H3K27me3-RADIP sample than in the Input sample (Figure 2C). We observed a significant decrease in RNA–DNA interactions at distances exceeding 1Mb in both Input and H3K27me3-RADIP samples. Upon comparing the lengths of RNA–DNA interactions from Input and H3K27me3-RADIP samples with topologically associating domains (TADs) derived from Hi-C data, it was evident that H3K27me3-RADIP significantly captures more long-range RNA–DNA interactions compared to Input. However, most interaction lengths were still shorter than those observed in the TADs from Hi-C. The average TAD length of approximately 0.8M indicates that DNA–DNA interactions are crucial in restricting how far the many RNAs can spread from their transcription sites to interact with distant DNA regions (Figure 2D). These results suggest that RNA–chromatin interactions mediated by H3K27me3 histone marks occurred at relatively long distances, with some RNAs interacting with chromatin separated by more than 100 kb from their transcribed sites. The RNA–DNA interaction matrix on chromosome 7 was enhanced as a representative region. On chromosome 7 in the H3K27me3-RADIP samples, for example, the interactions of *Kcnq1ot1* and *Tenm4* RNAs had longer distances between the RNA tags and their target DNA regions than in the Input samples (Figure 2E).

**Figure 2.**
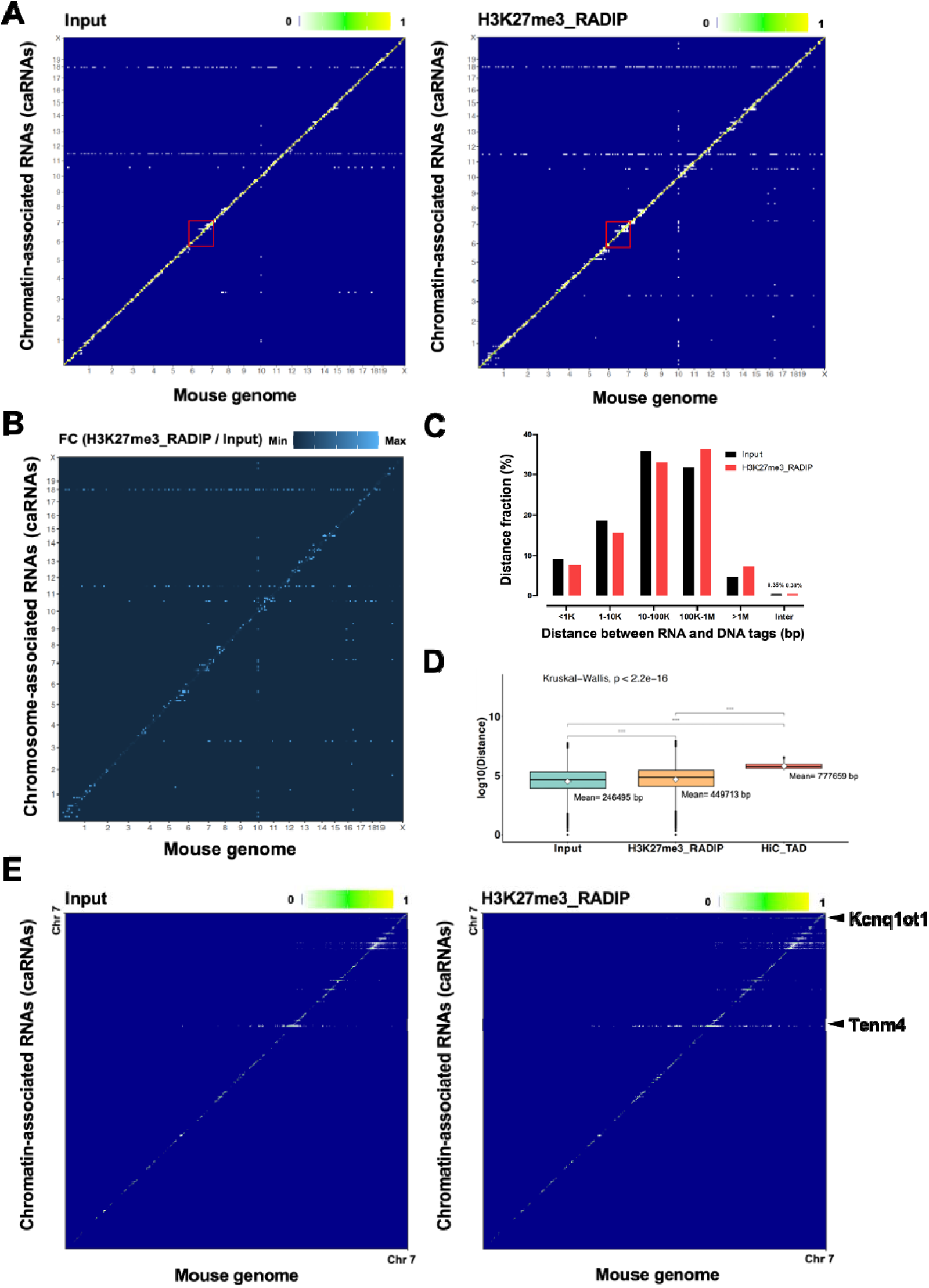
H3K27me3-RADIP enriched more long-range RNA–chromatin interactions compared to the Input. (**A**) Heatmap showing chromatin-associated RNAs across the whole genome in mouse ESCs. *Y*-axis: chromatin-associated RNAs (caRNAs). *X*-axis: mouse mm10 genome. Left: Input sample; right: H3K27me3-RADIP sample. (**B**) Calculation and visualization of the fold-change values after comparison of the H3K27me3-RADIP and the Input samples, showing that the H3K27me3-RADIP samples captured more long-range *cis* interactions and *trans* interactions that were far from the diagonal. FC, fold change. (**C**) RNA–chromatin interactions in Input (black) and H3K27me3-RADIP (red) samples, quantified according to the genomic distance between RNA and DNA tags. (**D**) The comparison of the lengths of RNA-DNA interactions from Input and H3K27me3-RADIP samples with TADs derived from Hi-C data showed that H3K27me3-RADIP significantly captures more long-range RNA-DNA interactions than the Input. However, most of these interaction lengths were still shorter than those observed in the TADs from Hi-C data. (**E**) The representative chromosome (chr.) 7 is enlarged, showing that some RNAs could interact with chromatin in a long-range *cis* manner and the interaction distance may be longer than that in the Input sample.

### Definition of RNA species enriched by H3K27me3-RADIP

We initially compared RNA tags captured by Input and H3K27me3-RADIP to confirm that RNA tags from H3K27me3-RADIP data could indeed interact with PRC2. By using the photoactivatable-ribonucleoside-enhanced cross-linking and immunoprecipitation (PAR-CLIP) dataset and the RNA immunoprecipitation (RIP-seq) dataset for EZH2 (a core component of PRC2) targeting and hence PRC2 targeting, we observed the number of RNA species harvested increased proportionately with sequencing depth in both libraries but was lower in the H3K27me3-RADIP library than in the Input library. Furthermore, the RNAs filtered out by H3K27me3-RADIP were predominantly transcribed from euchromatin regions not associated with H3K27me3 histone marks. This is evidenced by their lack of overlap with the EZH2-targeted PAR-CLIP and RIP-seq datasets, demonstrating that H3K27me3-RADIP technology effectively excluded some fraction of RNA–DNA interactions unrelated to PRC2 (Figure 3A). The substantial overlap of protein-coding RNA (pcRNA) species identified using H3K27me3-RADIP technology with those identified by EZH2-targeted PAR-CLIP and RIP-seq datasets, along with the notable overlap of long non-coding RNA (lncRNA) species identified by H3K27me3-RADIP technology with those found in the same EZH2-targeted PAR-CLIP and RIP-seq datasets, suggests that most H3K27me3-mediated RNA–DNA interactions are directly associated with PRC2 (Figure 3B).

**Figure 3.**
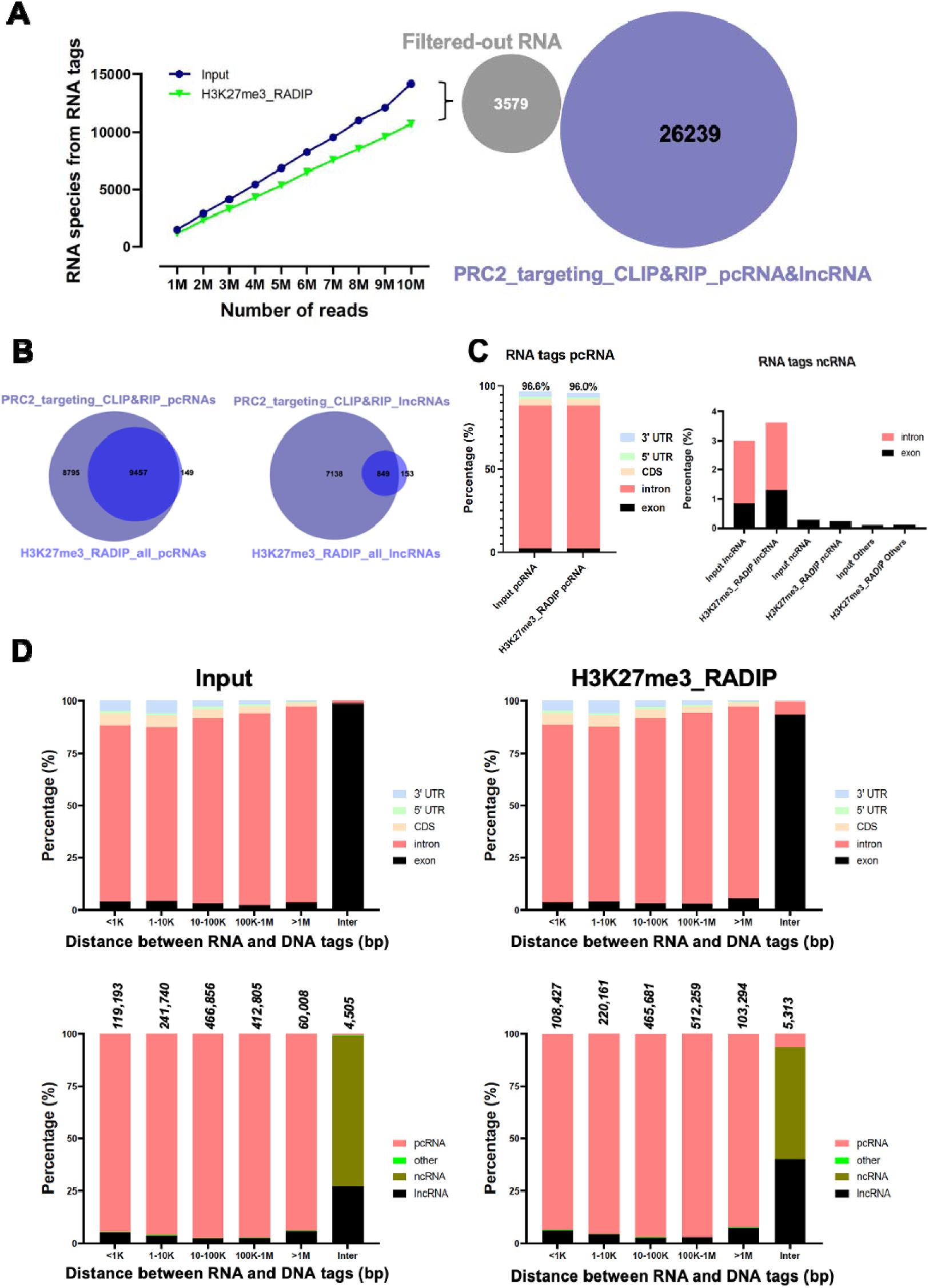
Characterization of RNA species in RNA–chromatin interactions mediated by H3K27me3 histone marks. (**A**) Evaluation of RNA species captured from Input and H3K27me3-RADIP samples in relation to sequencing depth. The number of RNA species harvested increased proportionately with sequencing depth in both libraries but was lower in the H3K27me3-RADIP library than in the Input library. Moreover, the filtered-out RNAs by H3K27me3-RADIP were predominantly transcribed from euchromatin regions not associated with H3K27me3 histone marks, as evidenced by their lack of overlap with the EZH2-targeted PAR-CLIP and RIP-seq datasets, demonstrating that H3K27me3-RADIP technology effectively excluded RNA–DNA interactions unrelated to PRC2. (**B**) Overlap of protein-coding RNA (pcRNA) species identified by H3K27me3-RADIP technology with those identified by the EZH2-targeted PAR-CLIP and RIP-seq datasets, as well as overlap of long non-coding RNA (lncRNA) species identified by H3K27me3-RADIP technology with those identified by the EZH2-targeted PAR-CLIP and RIP-seq datasets. (**C**) Origins of RNA tags. Both Input and H3K27me3-RADIP significant RNA tags were found to be dominated by pcRNA intronic regions (about 86%) (left columns). The origins of ncRNA are shown (right columns). (**D**) Input RNA–DNA interactions (left) and H3K27me3-RADIP RNA–DNA interactions (right) were quantified according to the genomic distance between the RNA and DNA tags. Both Input and H3K27me3-RADIP RNA–DNA interactions were catalogued and calculated according to the derived regions of the RNA tags and the classes of the RNA tags.

To investigate the RNA origins of RNA–DNA interaction pairs that participated in long-range interactions in the H3K27me3-RADIP samples, we analyzed the differences between the Input and H3K27me3-RADIP samples in the distributions of the significant RNA–DNA interaction pairs for various transcript biotypes and RNA regions. About 86% of the RNA–DNA interaction pairs in the Input and H3K27me3-RADIP samples were dominated by intronic RNA reads originating from protein-coding genes. In the H3K27me3-RADIP samples, the percentage of RNA–DNA interaction pairs from exonic RNA tags of lncRNAs was marginally increased (Figure 3C), although the increased tags largely corresponded to *Malat1* lncRNA. Our results agree with the previous studies that have highlighted abundant global binding of the *Malat1–*PRC2 complex on chromatin; this includes many *trans* interactions (55,56). The Input and the H3K27me3-RADIP samples displayed very similar profiles regarding RNA origins and interaction distances (Figure 3D). These results suggested that specific RNA origins or transcript biotypes may not contribute to long-range RNA–chromatin interactions mediated by H3K27me3.

Next, we investigated whether specific RNA species in the H3K27me3-RADIP samples reflected RNA–chromatin interactions dependent on H3K27me3 histone marks. We focused on transcripts from protein-coding genes (referred to as “pcRNAs”: Notably the 86% fraction of the pcRNAs consist of introns of protein-coding genes.) and lncRNA species captured by the Input and H3K27me3-RADIP samples. About 85% of the pcRNA species captured by H3K27me3-RADIP overlapped with those captured by Input. The numbers of pcRNA species captured by both samples were similar (Input: 9,288; H3K27me3-RADIP: 9,606). In contrast, only about 57% of the lncRNA species captured by H3K27me3-RADIP overlapped with those of Input, with H3K27me3-RADIP capturing more lncRNAs species than Input (Input: 852; H3K27me3-RADIP: 1,002) (Figure 4A). These results suggested that some lncRNAs were preferentially captured by the H3K27me3-RADIP technology and that these identified lncRNAs had greater potential to recruit PRC2, thereby facilitating the deposition of H3K27me3 histone modification marks. Among the shared pcRNAs and lncRNAs in the Input and H3K27me3-RADIP samples, about 10.5% of the pcRNAs and 53% of the lncRNAs were differentially enriched (including both Input- and H3K27me3-RADIP-enriched), as determined from the normalized counts of RNA–DNA interaction pairs (adjusted *P* ≤ 0.1 and absolute value of log2 fold change ≥ 0.5) (Figure 4B, top). The enriched pcRNAs and lncRNAs were queried by expression level by using the RNA-seq data. There were low levels of expression of pcRNAs and lncRNAs enriched in the H3K27me3-RADIP samples (Figure 4B, bottom).

**Figure 4.**
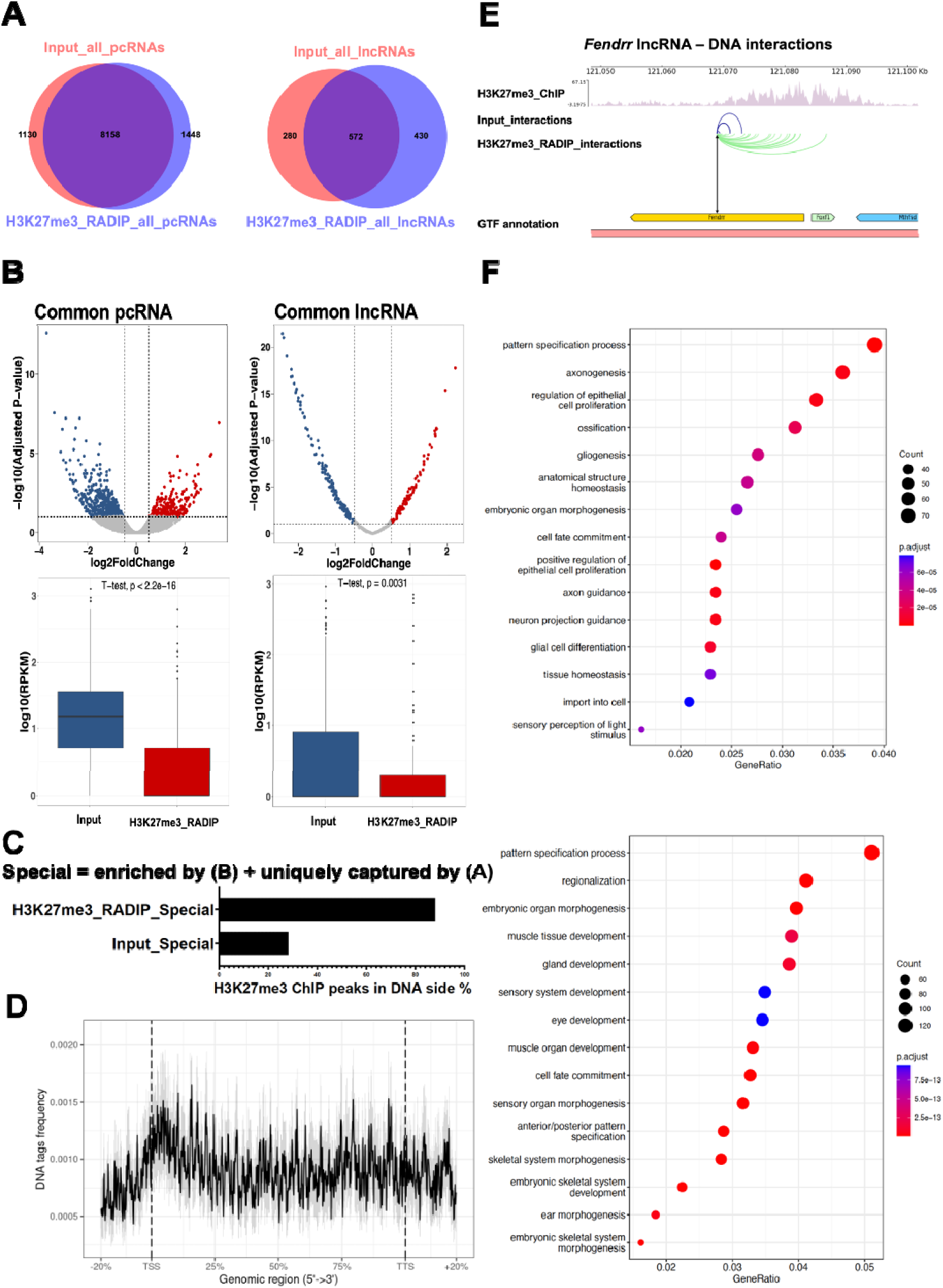
Definition of RNA species enriched by H3K27me3-RADIP. (**A**) Overlap of pcRNA species identified in Input and H3K27me3-RADIP samples, as well as overlap of long non-coding RNA (lncRNA) species identified in Input and H3K27me3-RADIP samples. (**B**) Left: differential interaction analysis of common pcRNAs detected in both Input and H3K27me3-RADIP samples (top). Expression of pcRNAs enriched in Input and H3K27me3-RADIP samples (bottom). Right: differential interaction analysis of common lncRNAs detected in both Input and H3K27me3-RADIP samples (top). Expression of lncRNAs enriched in Input and H3K27me3-RADIP samples (bottom). (**C**) Percentage of H3K27me3 ChIP peaks located in DNA-side tags from Input “Special” (enriched by differential interaction analysis of (**B**) + uniquely captured by (**A**)) RNA–chromatin interaction pairs and H3K27me3-RADIP “Special” (enriched by differential interaction analysis of (**B**) + uniquely captured by (**A**)) RNA–chromatin interaction pairs. (**D**) Distribution of DNA-side tags from H3K27me3-RADIP “Special” (enriched by differential interaction analysis of (**B**) + uniquely captured by (**A**)) RNA-chromatin interaction pairs across genomic regions. (**E**) Representative lncRNA–DNA interactions mediated by H3K27me3 histone marks (*Fendrr* lncRNA). Note that the source RNA region was set to the center of the *Fendrr* gene. (**F**) Gene Ontology term analysis of pcRNAs and lncRNAs from H3K27me3_RADIP “Special” RNAs (top), as well as target genes covered by the H3K27me3 ChIP peak regions of these RNAs (bottom).

In addition, the DNA targets of “Special” (enriched + unique) RNA species in the H3K27me3-RADIP samples (Supplementary Table S1) exhibited a substantially greater overlap than those in the Input samples with the H3K27me3 ChIP peaks (Figure 4C), along with a noticeable enrichment around transcription start sites (TSS) and promoter regions (Fig. 4D). These results suggest that the function of RNAs interacting with H3K27me3-marked genomic regions may be associated with the repressive roles of PRC2. These RNAs potentially recruit PRC2 to the promoter regions of genes, facilitating the deposition of H3K27me3 histone marks and ultimately silencing gene expression. For instance, the number of *Fendrr* lncRNA interactions was much higher in the H3K27me3-RADIP samples than in the Input samples. In addition, the target DNA tags of *Fendrr* lncRNA in the H3K27me3-RADIP samples were strongly correlated with the H3K27me3 ChIP peaks that covered the promoter region of adjacent *Foxf1* gene (Figure 4E). It has been reported previously that, in mouse E8.5 embryonic caudal ends, *Fendrr* lncRNA recruits PRC2 to the local genomic promoter region and represses expression of the adjacent *Foxf1* gene by depositing a strong H3K27me3 signal (57). We also presented the results for other lncRNAs, such as *A930004J17Rik*, *Zmiz1os1*, *A930037H05Rik*, and *Airn* lncRNA from “Special” RNA species in the H3K27me3-RADIP dataset as examples (Supplementary Figure S4 and Supplementary Figure S5A). We consistently observed that the RNA–DNA interaction numbers of these lncRNAs were substantially higher in the H3K27me3-RADIP samples than in the Input samples. Moreover, the target DNA tags of these lncRNAs in the H3K27me3-RADIP samples were strongly associated with the H3K27me3 ChIP peaks that covered the promoter regions of some genes, implying that these lncRNAs may participate in the repression of target gene expression through their collaboration with PRC2. Moreover, *Airn* lncRNAs have been reported to facilitate the spread of PRCs across distinct chromatin domains (58,59). Our H3K27me3-RADIP also showed that nearly all the *Airn* lncRNA–DNA interactions were mediated by *Airn* intron-containing transcripts (Supplementary Figure S5B). To determine whether the *Airn* introns captured by H3K27me3-RADIP were actually associated with chromatin, we isolated the chromatin fraction from mESCs both with and without treatment with GSK126 (an EZH2 inhibitor (60)). This was followed by reverse transcription–quantitative polymerase chain reaction (RT-qPCR). We designed PCR primer sets for the intronic region of *Airn* lncRNA and the exonic region of *Nanog* mRNA. The percentage of *Airn* introns in the chromatin fraction was significantly reduced after treatment with GSK126, an EZH2 inhibitor, but the percentage of *Nanog* mRNA exons used as a negative control remained unchanged (Supplementary Figure S5C and see Supplementary Methods). These results implied that “Special” RNA species identified in the H3K27me3-RADIP samples were physically associated with the chromatin in a manner that was dependent either on the catalytic activity of EZH2 or on H3K27me3 histone marks. However, we do not know whether those intronic RNA tags are intron fragments, pre-mature mRNAs, or nascent RNAs. Finally, we performed a gene ontology (GO) term analysis of pcRNAs and lncRNAs from H3K27me3-RADIP “Special” RNAs (Figure 4F, top), as well as of DNA target genes covered by the H3K27me3 ChIP-seq peak regions of these H3K27me3-RADIP “Special” RNAs (Figure 4F, bottom). We found that they were indeed involved in some expected biological processes, such as cell development, cell proliferation, cell differentiation, and cell fate commitment. This was in agreement with the partial GO term results for PRC2 in mESCs (61), not only for the DNA target genes but also for the “Special” RNAs. These results suggested that H3K27me3-associated RNAs may participate in the establishment of repressive marks for their target genes.

### Relationship between 3D chromatin structures and H3K27me3-mediated RNA–DNA interactions

Based on our previous RADICL-seq data, we noticed that the DNA tags of RNA–chromatin interactions are significantly enriched at TAD boundaries (12). These TAD boundaries serve to constrain the spread of transcriptional activities, acting as barriers that prevent the free diffusion of RNA migration. This suggests that RNA–DNA interactions may, to some extent, reflect genomic interactions, which are closely related to RNA production and regulation. To further explore this, we utilized H3K27me3-RADIP captured long-range RNA–DNA interactions to characterize TADs associated with the activity of PRC2. This approach has the potential to reveal new insights into the structural and regulatory mechanisms governing chromatin remodeling and gene expression. Initially, we analyzed the hierarchical domains mediated by RNA–chromatin interactions using an approach analogous to the topologically associating domain (TAD) approach utilized in the Hi-C technology (62). We identified the hierarchical domains of RNA–DNA interaction pairs in Input and H3K27me3-RADIP and referred to them as “TAD-like domains” (TLDs) (49). We illustrated the TLDs on chromosome 6 as an example (Figure 5A). The TLDs were highly correlated across the Input and H3K27me3-RADIP samples on chromosome 6 (Pearson’s correlation coefficient = 0.7). After comparing the Input samples with the H3K27me3-RADIP samples at the level of the TLDs, we defined anchors and collected the anchor information of TLDs that had changed between Input and H3K27me3-RADIP on a genome-wide level. Subsequently, we determined which RNA species were related to the changed TLDs; these were referred to as “Changed TLD” RNAs. We summarized the details of the changed TLDs on chromosome 6 (Supplementary Figure S6A), then, listed the “Common TLD” RNAs and “Changed TLD” RNAs on chromosome 6 (Supplementary Figure S6B). We extracted 853 RNA species in total and defined them as “Changed TLD” RNAs on a genome-wide scale. The “Changed TLD” RNAs were likely to have contributed to the gain or loss of anchors. We then intersected the “Changed TLD” RNAs with the H3K27me3-RADIP “Special” RNAs and found that 20.9% (178 of 853) of the RNAs in the two categories overlapped with each other. In contrast, 8.4% (72 of 853) of the “Changed TLD” RNAs overlapped with the Input “Special” RNAs. The overlapping section between “Changed TLD” RNAs and H3K27me3-RADIP “Special” RNAs was referred to as the “Overlap” RNAs (Supplementary Table S2). These “Overlap” RNAs were markedly enriched in TLDs gained in the H3K27me3-RADIP samples and were distinct from those in the Input samples in terms of the quantity and interaction pattern of RNA–DNA interactions. These “Overlap” RNAs potentially interact with chromatin along with PRC2, leading to a gradual compaction of target regions after the deposition of H3K27me3 histone marks, and ultimately contributing to the formation of TADs. This specific association highlights the role of these RNAs interacting with H3K27me3-marked genomic regions. (Figure 5B). To confirm that the RNA–DNA interactions associated with the “Overlap” RNAs are more closely correlated with TADs, we retrieved publicly available Hi-C data on mESCs (63) and examined the relationship between DNA–DNA interaction pairs and RNA–DNA interaction pairs. The frequencies of normalized RNA–DNA contacts for *cis* interactions in the Input “Special” dataset and the H3K27me3-RADIP “Special” dataset were both modestly correlated with the normalized Hi-C DNA–DNA contacts at the same 25-kb bin resolution (Pearson’s correlation coefficient = ∼ 0.48) (Figure 5C); this was consistent with our previous observations (12). To analyze the fraction highly enriched in H3K27me3-RADIP-gained TLDs, we then focused on the “Overlap” dataset. The RNA–DNA interaction pairs from the “Overlap” dataset were even more similar to the Hi-C DNA–DNA contacts (Pearson’s correlation coefficient = 0.65), and the DNA targets were enriched in the binding motif of CCCTC-binding factor (CTCF) (Figure 5D), indicating that the “Overlap” RNAs may play a role in the establishment and maintenance of H3K27me3-mediated 3D chromatin structures through the activity of PRC2. These findings suggest a dynamic interplay where “Overlap” RNAs may act as guides or stabilizers in the formation of chromatin domains through long-range RNA–DNA interactions that extend beyond immediate neighboring regions (Figure 5E).

**Figure 5.**
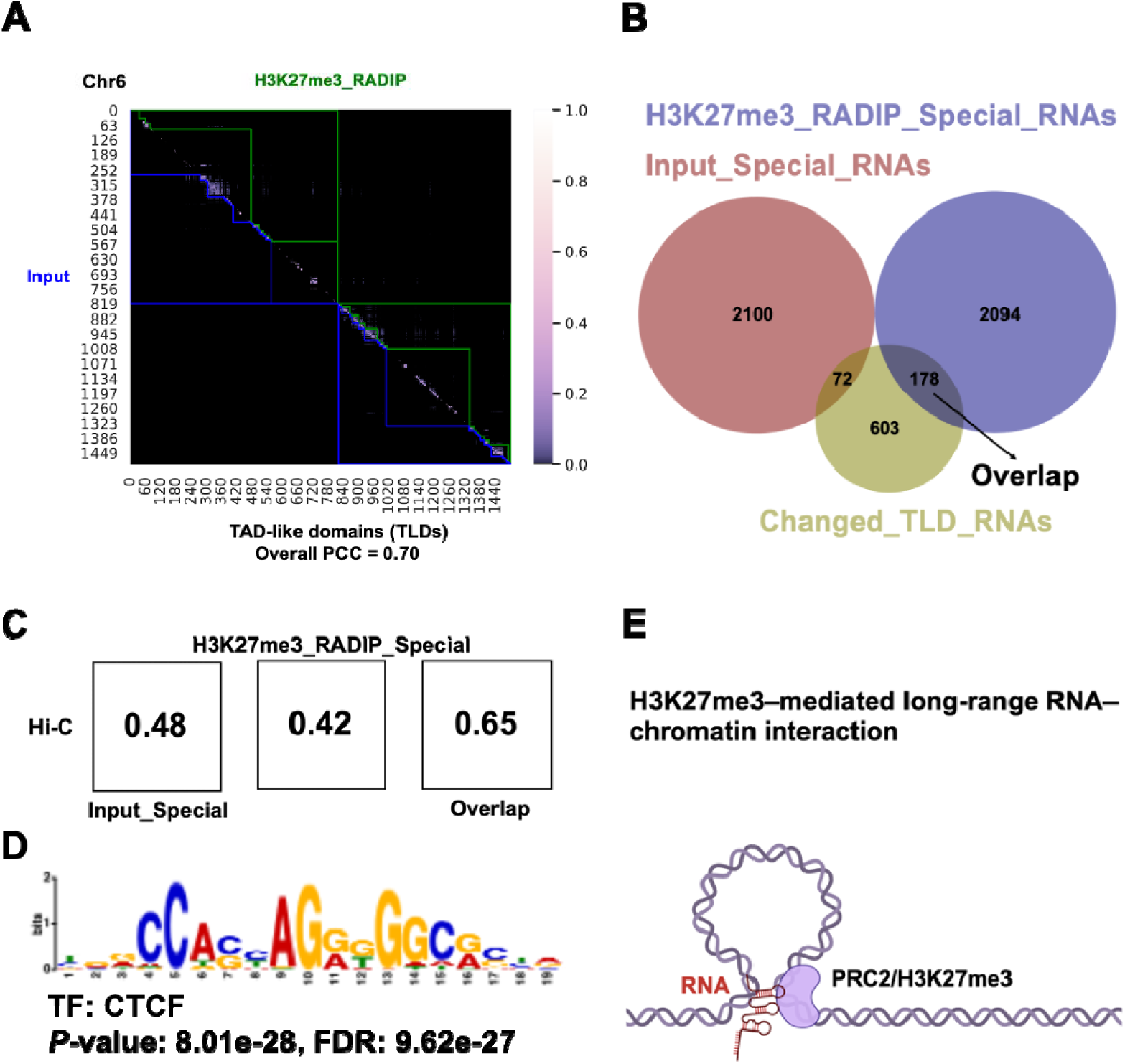
Identification of RNAs that may contribute to the formation of PRC2-associated TADs. (**A**) Comparison of TAD-like domains (TLDs) on the basis of Input and H3K27me3-RADIP samples. Chromosome (chr.) 6 is presented as an example. (**B**) Overlap of RNA species among the “Changed TLD” RNAs, Input “Special” RNAs, and H3K27me3_RADIP “Special” RNAs. (**C**) Pearson’s correlation coefficients between RNA–DNA interactions and DNA–DNA interactions. (**D**) The DNA targets of “Overlap” RNAs were enriched in the binding motif of CCCTC-binding factor (CTCF). (**E**) The essential role of “Overlap” RNAs in forming PRC2-associated TADs, which facilitate specific long-range RNA–DNA interactions.

### Identification of lineage-specific RNA-mediated transcriptional silencing

To investigate lineage-specific RNA-mediated transcriptional silencing, we utilized the publicly available data of a differentiation model, as follows. This model involves the culture of mESCs under conditions with leukemia inhibitory factor (LIF) and induced differentiation into embryoid bodies following the removal of LIF and supplementation with 10 µM retinoic acid (RA) for 6 days (64). To elucidate the RNA-induced genomic distribution of H3K27me3 and its impact on gene expression, RNA-seq and ChIP-seq data were downloaded at the day 0 and 6 time points. Differential expression analysis of RNA-seq data revealed significant gene expression changes between the pluripotent and differentiated states, as evidenced by adjusted P-values ≤ 0.05 and absolute value of log2 fold change ≥ 1.5 (Figure 6A). Subsequent differential intensity analysis of H3K27me3 ChIP peak regions identified significantly changed genomic regions between the two stages (adjusted P ≤ 0.05 and absolute value of log2 fold change ≥ 0) (Figure 6B). Further analysis correlated gene expression changes with fluctuations in H3K27me3 intensity. By overlapping the datasets, we identified a subset of genes that, upon differentiation, exhibited a decrease in H3K27me3 intensity and a corresponding increase in expression. This reinforces the role of H3K27me3 as repressive histone marks and underscores the crucial role of these marks in maintaining the silencing of these differentiation-related genes in mESCs (Figure 6C). GO term analysis also confirmed that these genes are significantly associated with biological processes including cell development, differentiation, and fate commitment (Figure 6D). Subsequently, we identified RNAs that overlapped with these differentiation-related genes from H3K27me3-RADIP data in mESCs. Among the H3K27me3-RADIP “Special” RNAs, we found that 10% of the H3K27me3-RADIP “Special” lncRNAs (55 of 534) and 15% of the H3K27me3-RADIP “Special pcRNAs (265 of 1,738) were interacting with these differentiation-related genes.

**Figure 6.**
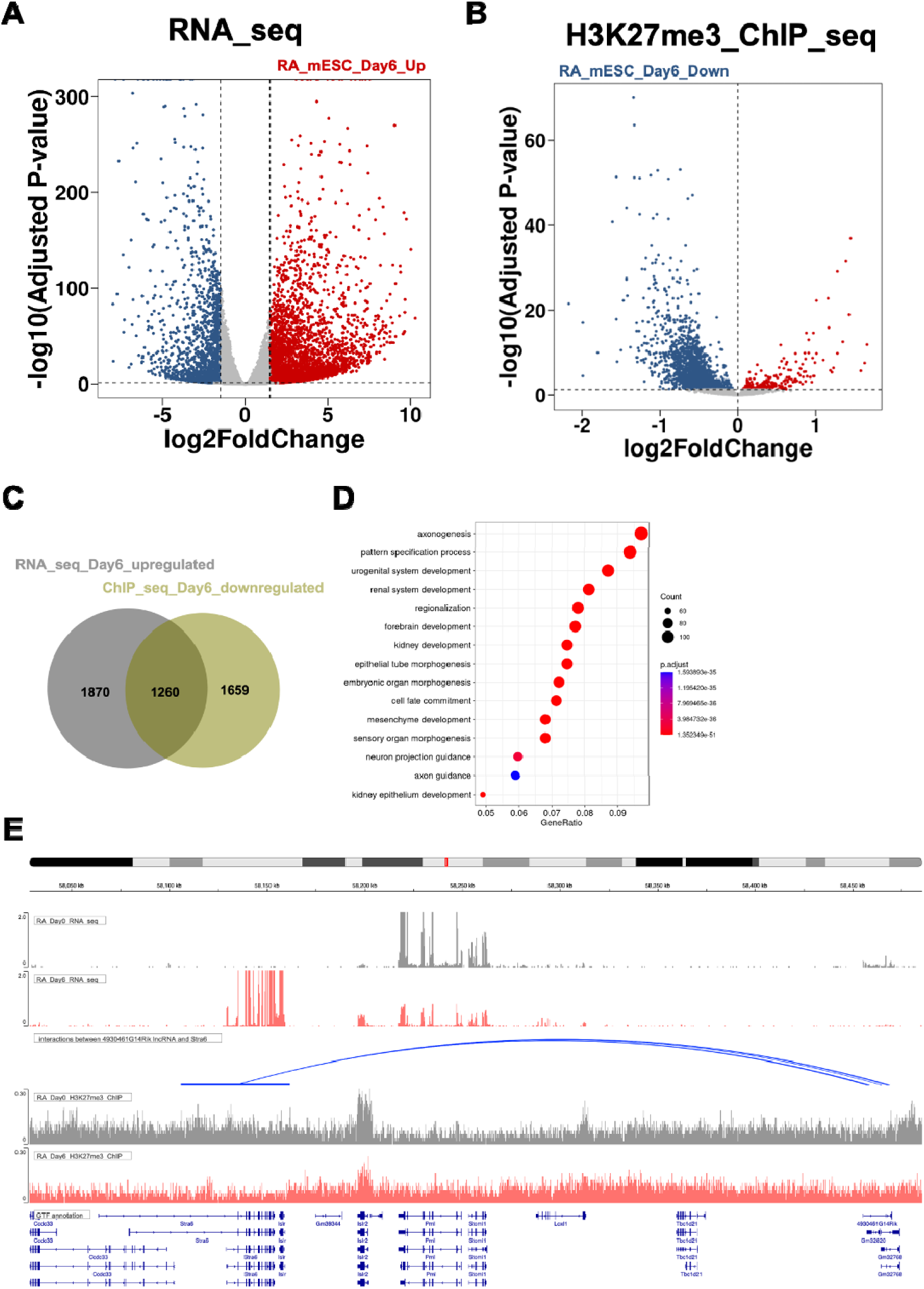
Definition of RNA-mediated lineage-specific transcriptional silencing. (**A**) Differential expression analysis of RNA-seq data revealed significant gene expression changes between the pluripotent and differentiated states (adjusted P-values ≤ 0.05 and absolute value of log2 fold change ≥ 1.5). (**B**) Differential intensity analysis of H3K27me3 ChIP peak regions identified out significantly changed genomic regions between the two states (adjusted P ≤ 0.05 and absolute value of log2 fold change ≥ 0). (**C**) Overlap of significantly changed RNA-seq data and ChIP-seq data identified a subset of genes that, upon differentiation, showed a decrease in H3K27me3 intensity and a corresponding increase in expression. (**D**) GO term analysis of these identified genes. (**E**) Visualization of interactions between *4930461G14Rik* lncRNA and the *Stra6* gene. the *4930461G14Rik* lncRNA guides PRC2 to the TSS of the *Stra6* gene, facilitating the deposition of H3K27me3 and thereby silencing the *Stra6* gene in mESCs. As differentiation progresses, the expression of *4930461G14Rik* lncRNA ceases, leading to diminished guidance, reduced PRC2 localization, and a consequent decrease in H3K27me3 intensity at the TSS of *Stra6* gene. These changes result in a significant up-regulation of *Stra6* gene expression, which is critical for retinol uptake and thus promotes cell differentiation.

For example, we visualized the interactions between the *4930461G14Rik* lncRNA and the *Stra6* gene. Initially, an H3K27me3 ChIP peak region was identified covering the transcription start site (TSS) of the *Stra6* gene in mESCs. Upon differentiation, a significant decrease in H3K27me3 intensity at this region was observed. From the H3K27me3-RADIP data, we found that *4930461G14Rik* lncRNA specifically interacts with this region in mESCs. This suggests that the *4930461G14Rik* lncRNA may guide PRC2 to the TSS of the *Stra6* gene, facilitating the deposition of H3K27me3 and thereby silencing the *Stra6* gene in mESCs. We postulated that as the process of spontaneous differentiation progresses, the expression of *4930461G14Rik* lncRNA ceases, resulting in diminished guidance, reduced PRC2 localization, and a consequent decrease in H3K27me3 intensity at the TSS of the *Stra6* gene (Figure E). These changes may result in a significant up-regulation of *Stra6* gene expression.

### RNA motifs contributing to chromatin associations

We further investigated whether the motifs in the RNA sequences were involved in facilitating RNA interactions with the genomic regions mediated by H3K27me3 histone marks (See Materials and methods). We identified 67 motifs with a *P*-value threshold of 0.05 distributed across the RNA tags from the RNA–DNA interaction pairs in the H3K27me3-RADIP samples (Supplementary Figure S7 to S10). We selected four representative motifs (Figure 7A; see Materials and Methods). Among them, two left motifs, (GGGG)n and (GGAA)n, were G-rich, as has been previously reported among PRC2-binding motifs (22). Another motif was C- and U-rich, which was consistent with a previous analysis of EZH2- or SUZ12-binding motifs (65). The last motif was A- and U-rich and is indicative of AU-rich elements. These findings indicated that many RNA-binding proteins (RBPs) were involved in these H3K27me3-mediated RNA–chromatin interactions (66–69).

**Figure 7.**
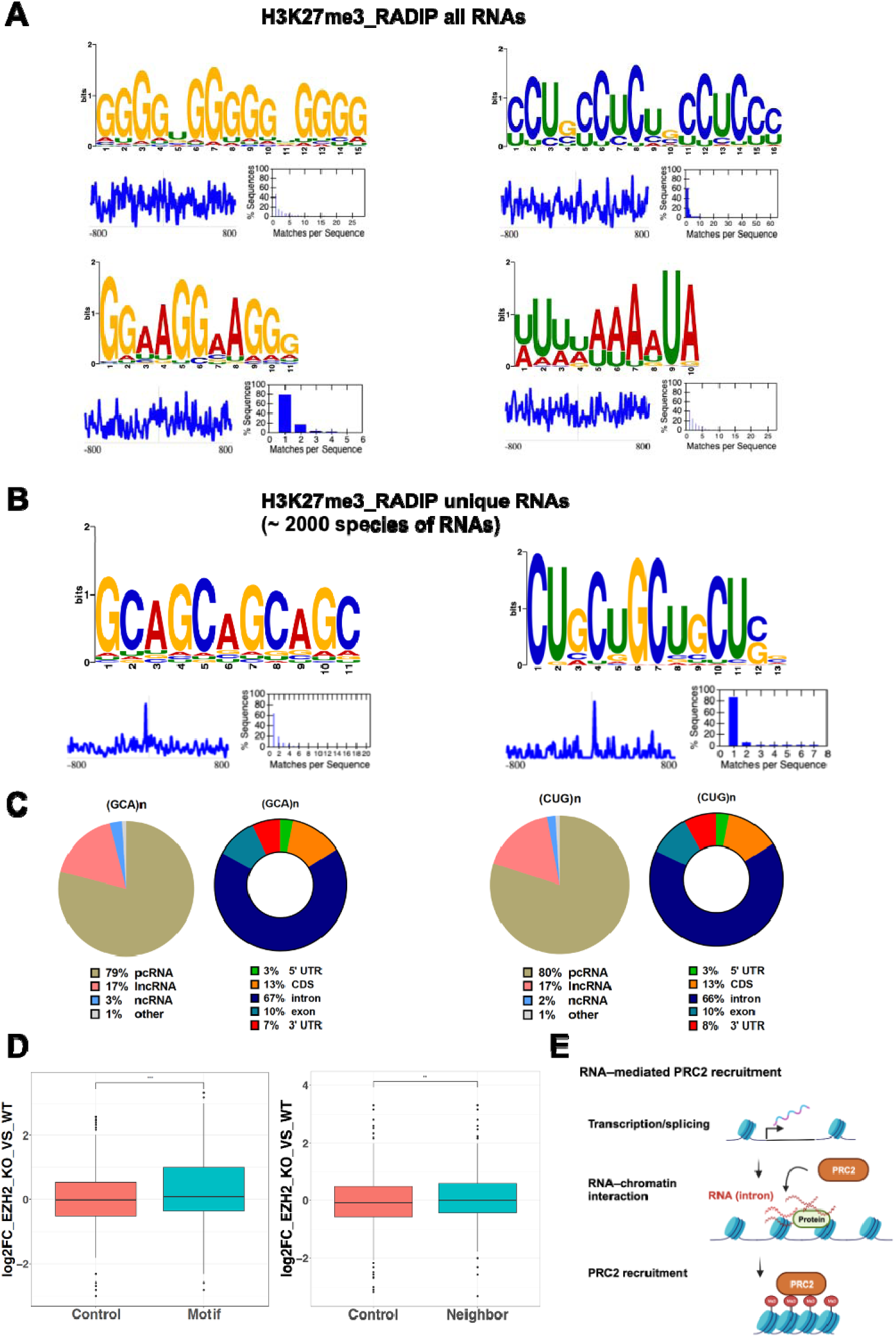
Motif enrichment analysis in RNA tags of H3K27me3-RADIP samples. (**A**) Representative sequence logos of motifs enriched in all RNA tags from H3K27me3-RADIP RNA–chromatin interaction pairs. RNA tags were extended ± 800 nt from the center points. (**B**) Highly enriched motifs based on the RNA tags from unique RNA species detected only in the H3K27me3-RADIP samples and not in the Input samples. RNA tags were extended ± 800 nt from the center points. Note that the GCA motif was found in 87% of the queried RNA tags and the CUG motif was found in 85% of the queried RNA tags. (**C**) Distributions of RNA classes and RNA regions, based on RNA tags containing the GCA motif and CUG motif. (**D**) Left: Expression-level changes of motif-containing RNAs and the same number of randomly selected control genes from the mouse mm10 genome after EZH2 knockout in mESCs. Right: Expression-level changes of neighboring genes (± 3 kb from the motif-containing RNAs) and the same number of randomly selected control genes from the mouse mm10 genome after EZH2 knockout in mESCs, suggesting that motif-containing RNAs may couple with PRC2 to repress gene expression in the whole neighborhood of the PRC2-binding sites via the deposition of H3K27me3 histone modification marks. (**E**) After RNAs targeted by PRC2 were transcribed and spliced, their intronic regions are attached to chromatin directly or via proteins to help recruit PRC2 to chromatin and deposit H3K27me3 histone marks (Me3).

To identify more specific motifs associated with H3K27me3-mediated RNA–chromatin interactions, we focused on RNA species unique to H3K27me3-RADIP (1906 species). A total of 69 motifs were enriched in the unique RNA species, with a *P*-value threshold of 0.05 (Supplementary Figure S11 to S14), and about 20 of these motifs overlapped with, or were similar to, the motifs that were identified across all RNAs in the H3K27me3-RADIP samples. From the non-overlapping and non-similar motifs, we selected two representative motifs (Figure 7B and see Materials and methods), (GCA)n and (CUG)n. We examined the source RNAs of the two motifs and found that the two motifs were mostly mapped to pcRNAs (about 79%). In addition, we observed that the motifs located in the intronic regions accounted for 66% to 67% of all the RNA regions (Figure 7C). To investigate the potential roles of RNAs containing GCA repeat and CUG repeat motifs, we investigated their links with PRC2. We retrieved the RNA-seq data for the conditional knockout of EZH2 and examined the expression-level changes of their source genes (i.e., the genes that were transcribed into RNAs containing GCA repeat and CUG repeat motifs). The expression levels of the motif-containing RNAs were significantly increased (*P*-value = 6.4e-10) after EZH2 knockout compared with the levels of the control RNAs (randomly selected and numerically equivalent to comparison RNAs), suggesting that RNAs containing these motifs contribute to the guidance of PRC2 to genomic regions, leading to the massive accumulation of H3K27me3 histone marks and gene repression (Figure 7D, left). We also examined the expression-level changes of the neighboring genes (± 3 kb) around the source gene after EZH2 knockout and found that these levels were significantly elevated (*P*-value = 0.0033) as well (Figure 7D, right). These results suggested that GCA-repeat- and CUG-repeat-motif-containing RNAs couple with PRC2 to repress gene expression in the entire neighborhood of the PRC2-binding sites by depositing H3K27me3 histone marks.

Taken together, these data indicated that the intronic regions of protein-coding genes associated with chromatin may act via intermediary proteins and assist PRC2 in depositing H3K27me3 histone marks locally or in spreading H3K27me3 histone marks in the surrounding regions (Figure 7E).

## DISCUSSION

We developed RADIP, a proximity-ligation-based method that combines RADICL-seq with immunoprecipitation with antibodies against specific proteins. We evaluated RADIP’s applicability by using an antibody against H3K27me3 histone marks associated with repressive chromatin regions. We identified a spectrum of caRNAs enriched at and around PRC2-binding chromatin sites and H3K27me3 peak regions in mESCs; these caRNAs may be involved in PRC2-mediated gene repression for pluripotency of mESCs and the establishment of PRC2-mediated three-dimensional genome architecture.

RADIP retains the primary advantages of RADICL-seq over other methods. Firstly, chromatin digestion is carried out using DNase I, resulting in a higher resolution compared to digestion with restriction enzymes. The distribution of cut sites for restriction enzymes is often uneven, which can lead to fewer detectable RNA interaction genomic regions and subsequently result in low DNA coverage. The sequence-independent digestion of chromatin by DNase I enables RADIP to overcome these resolution limitations, enhancing the ability to identify RNA-bound regions across the genome and providing a more comprehensive view of RNA–DNA interactions, irrespective of restriction site distribution. Secondly, the use of EcoP15I to generate RNA and DNA reads of uniform size improves unique alignment to the genome (Supplementary Figure S2A). Thus, we increase our ability to detect RNA–DNA interactions that are uniquely mapped. Notably, when comparing the percentage of sequencing reads that can be mapped to the mouse genome on both DNA and RNA sides, RADIP outperforms RedChIP, with more than a four-fold increase in the percentage of unique mapping reads (54% versus 13%). Additionally, we utilized the CHiCANE package (40), which was originally designed for the analysis of CHi-C data analysis, to statistically evaluate RNA–DNA interactions in RADIP data. We computationally eliminated background RNA–DNA interactions, which include a significant number of *trans* interactions.

By using H3K27me3-RADIP, we identified specific RNAs that were associated with H3K27me3 histone marks and were expressed at low levels but interacted with chromatin at longer distances from transcribed sites compared to the Input sample. Notably, a majority of the RNA tags originated from the intronic regions of protein-coding genes, implying a potential role for intronic pcRNAs as lncRNAs, although we do not yet know whether these intronic RNA tags are intron fragments, pre-mature mRNAs, or nascent RNAs. Motif enrichment analysis of RNAs captured by H3K27me3-RADIP revealed several enriched motifs, including those known to be targeted by EZH2 or PRC2. Among these, we identified two specific ones—GCA repeat and CUG repeat motifs—located predominantly in the intronic regions of protein-coding genes. By using the RBPDB database (70), we performed a subsequent analysis of the RBPs potentially interacting with these motifs and identified several splicing factors, including ‘splicing factor, arginine/serine-rich 9’ (SFRS9), muscleblind-like splicing regulator 1 (MBNL1), and Y box binding protein 1 (YBX1). YBX1 is a nucleic-acid-binding protein that participates in various DNA- or RNA-dependent processes, including mRNA translation, splicing, and transcription, and DNA repair (71). YBX1 overlaps highly with PRC2 binding genome-wide; it controls PRC2 distribution and decreases H3K27me3 levels, suggesting that it fine-tunes PRC2 activity to regulate spatiotemporal gene expression in embryonic neural development (72), although it is unclear whether or not RNA binding to YBX1 plays important roles in these processes. Our results demonstrated that RNAs with the GCA repeat and CUG repeat motifs were highly repressed by PRC2, unlike control RNAs (Figure 7D). This suggests that YBX1 may compete with other proteins binding to these RNAs with GCA repeat and CUG repeat motifs, highlighting its potential role in fine-tuning PRC2 activity and helping to regulate gene expression.

Although our RADIP technology and data will act as valuable resources for the study of caRNAs, there is still room for improvement. One limitation of RADIP is the relatively large number of cells required (about 20 million). This requirement largely depends on the number of target protein molecules present and the efficacy of the antibodies against these target proteins. Optimizing these factors could potentially reduce the cell number needed and enhance the overall feasibility of the RADIP technology. In another study, Khyzha *et al.* described RT&Tag (Reverse Transcribe and Tagment) as a proximity-labeling tool that facilitates the mapping of chromatin-associated RNAs (73). RNAs associated with a specific chromatin epitope are targeted by a specific antibody followed by a protein A-Tn5 transposome. Then, localized reverse transcription generates RNA–cDNA hybrids that are tagmented by Tn5 transposases for downstream next-generation sequencing. RT&Tag requires 0.1 million cells per sample, although Khyzha *et al.* tested only *Drosophila* S2 cells. They used antibodies to target the RNA-associated dosage compensation complex (MSL2), H3K27me3, and post-transcriptionally modified RNAs with m6A (N6-methyladenosine).

The RADIP method can be potentially used to identify RNAs associated with genomic regions occupied by any protein of interest. Although here we used an anti-H3K27me3 antibody to demonstrate how the RADIP method was applicable to the study of RNAs associated with repressive histone marks, we could use any antibodies against target proteins—for example, antibodies against transcription factors, CTCF, and RNA modifications. RADIP relies on proteins with known functions. Future advancements may focus on expanding the applicability of RADIP to *de novo* discoveries. Although the ENCODE Consortium has used eCLIP to generate the largest coherent public set of CLIP data, covering 150 RNA-binding proteins in two cell types (HepG2 and K562) (74), we still lack a complete catalog of RBPs in other cell types, as discussed in another study (73). In combination with RBP information, the RADIP method is likely to enable us to identify new functions for caRNAs.

In summary, RADIP technology will enable us to accelerate the exploration of a variety of RNA chromatin-regulatory activities mediated by specific proteins and determine the functions of new classes of RNAs.

## DATA AVAILABILITY

The RADIP data have been deposited in the GEO database under accession number GSE260447.

## SUPPLEMENTARY DATA

1. Supplementary Figures
2. Supplementary Methods
3. Supplementary Table S1-2

## ACKNOWLEDGEMENTS

X.S. and M.K. designed and conducted the experiments, X.S. performed the informatics analysis, S.T. performed the wet lab experiment, and Y.S. and P.C. supervised the study. X.S., M.K., and P.C. wrote the manuscript. We thank Dr. Lokesh Tripathi for comments on the manuscript.

