## Supplementary figures for "RADIP technology comprehensively identifies H3K27me3-mediated RNA-chromatin interactions"

#### Figure S1.

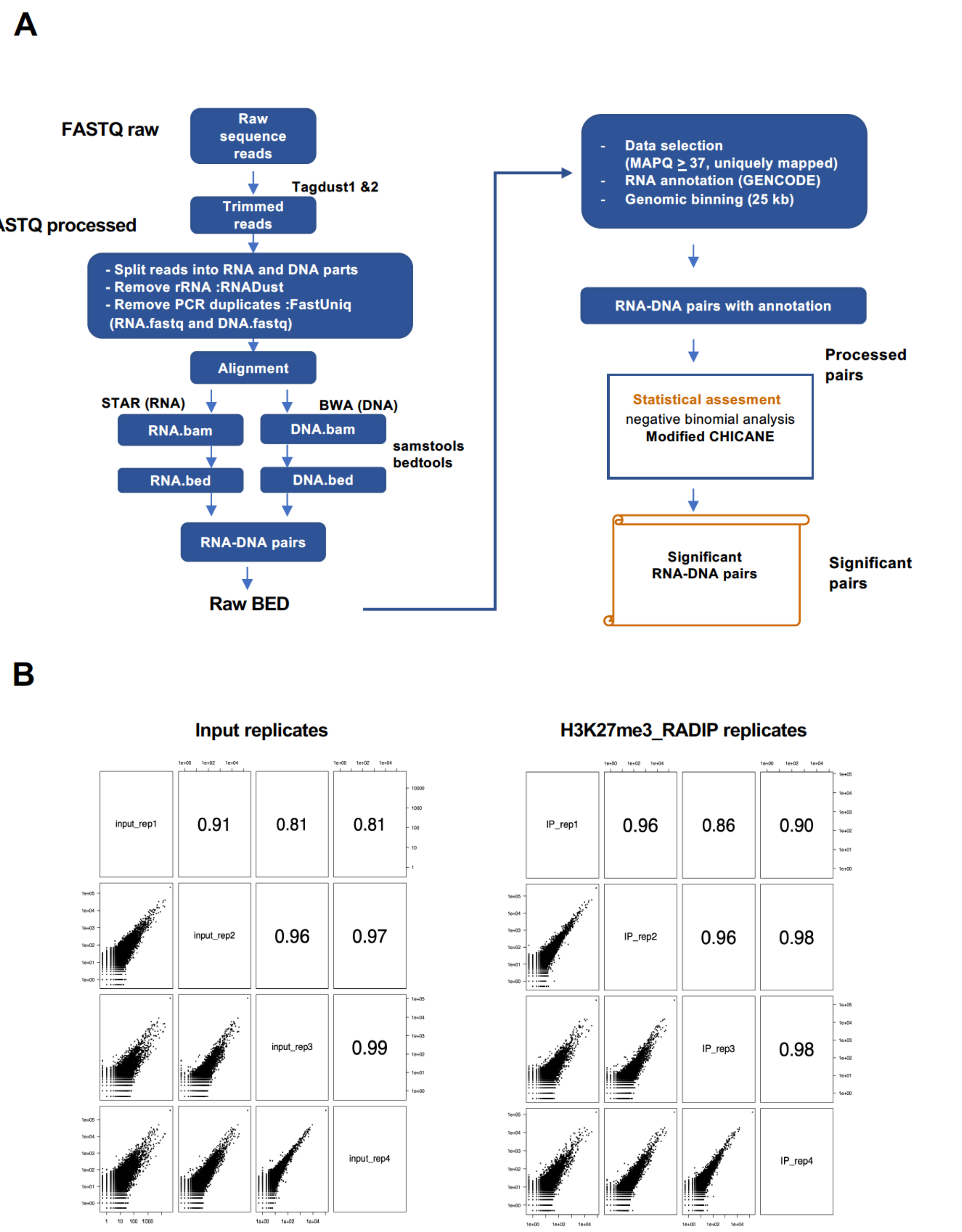


**Figure S1. Data-processing flowchart and high reproducibility of H3K27me3_RADIP technology.**

(**A**) Starting with the raw sequence reads, the adaptor sequences were trimmed off and the flanking sequences were split into the respective mate pairs of RNA and DNA reads on the basis of the orientation of the internal bridge adaptor. After the rRNA had been removed, data filtering and genomic alignment were performed. Only uniquely mapped RNA–DNA pairs were stored and annotated. The genome-wide RNA–DNA interactome was calculated in a matrix by evaluating captured RNA–DNA pairs between RNA species and genomic bins of 25 kb, with reporting of the processed pairs of all chromatin-associated RNAs on the binned genome. RNA–DNA pairs were processed by background removal, and significant pairs were output for downstream analysis. (**B**) Following these methodologies, RADIP technology exhibited high reproducibility among four Input replicates and four H3K27me3-RADIP replicates. BAW, Burrows–Wheeler Aligner; MAPQ, mapping quality.

#### Figure S2.

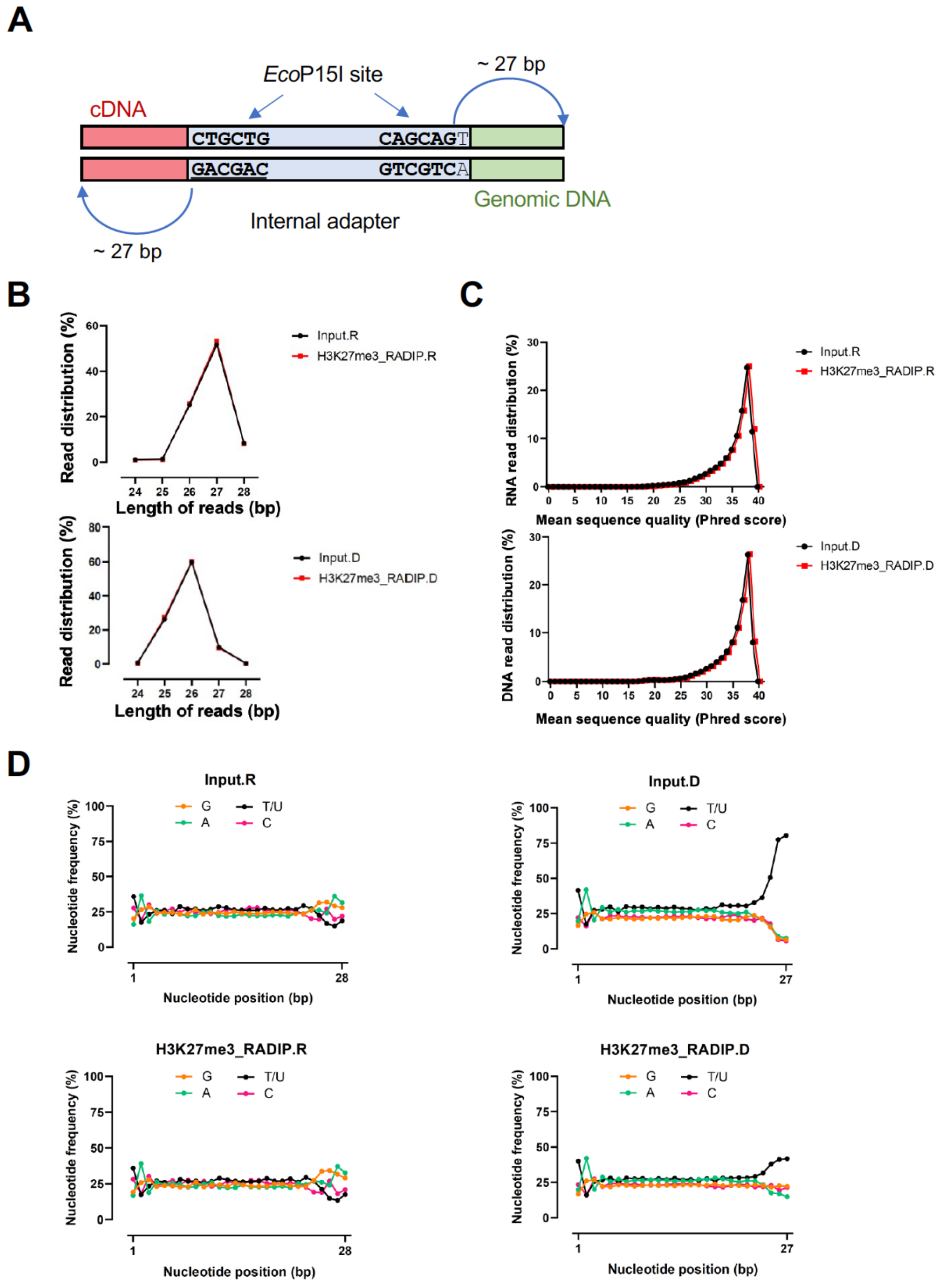


**Figure S2. H3K27me3_RADIP data quality and features.**

(**A**) Features of RNA–adaptor–DNA chimeric molecules. (**B**) Summary of lengths of RNA and DNA tags of Input and H3K27me3-RADIP samples. (**C**) Report of the base-call accuracy of RNA and DNA tags of Input and H3K27me3-RADIP samples. (**D**) Nucleotide frequencies of RNA (left) and DNA (right) tags of Input and H3K27me3-RADIP samples, showing that the distributions of nucleotides in the RNA and DNA tags were uniform, with the exception of the overhang of the T and Ecop151 recognition motif on the two ends of the DNA tags.

#### Figure S3.

A

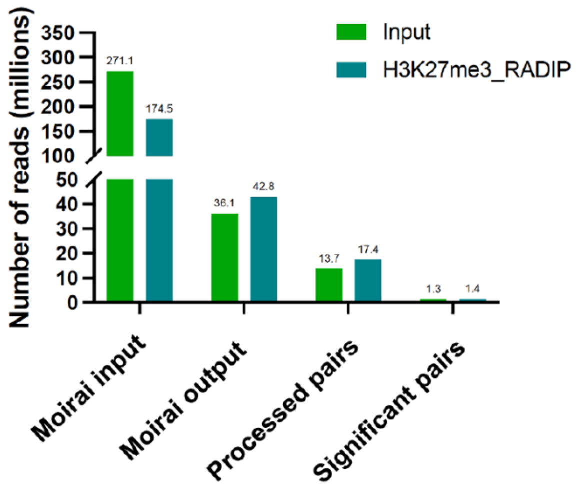


B

**Processed pairs**

| Before (RNA-DNA pairs) | *Cis* reads number | *Trans* reads number |
|---|---|---|
| Input | 4,269,577 | 9,429,349 |
| H3K27me3_RADIP | 4,684,311 | 12,675,142 |

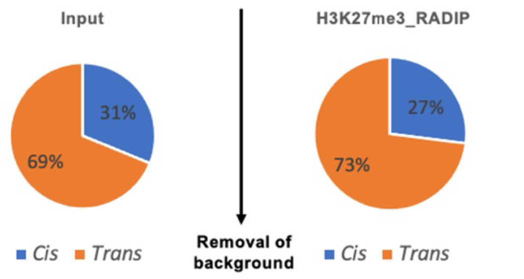


**Significant pairs**

| After (caRNA-DNA pairs) | *Cis* reads number | *Trans* reads number |
|---|---|---|
| Input | 1,300,602 | 4,505 |
| H3K27me3_RADIP | 1,409,822 | 5,313 |

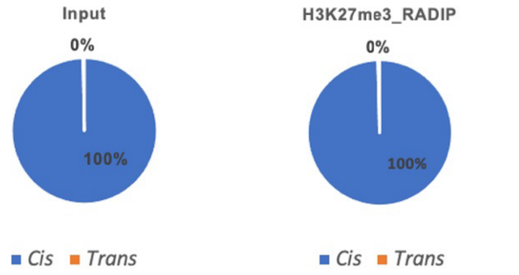


**Figure S3. Details of background removal.**

(**A**) Summary of sequenced Input and H3K27me3-RADIP libraries. Numbers above the bars represent the replicate-combined numbers of raw reads or the mate pairs remaining after the individual data-processing steps. (**B**) After removal of the background, many of the *trans* interactions were eliminated owing to their low frequency. The proportions and actual numbers of *cis* and *trans* RNA–DNA interaction pairs are shown.

#### Figure S4.

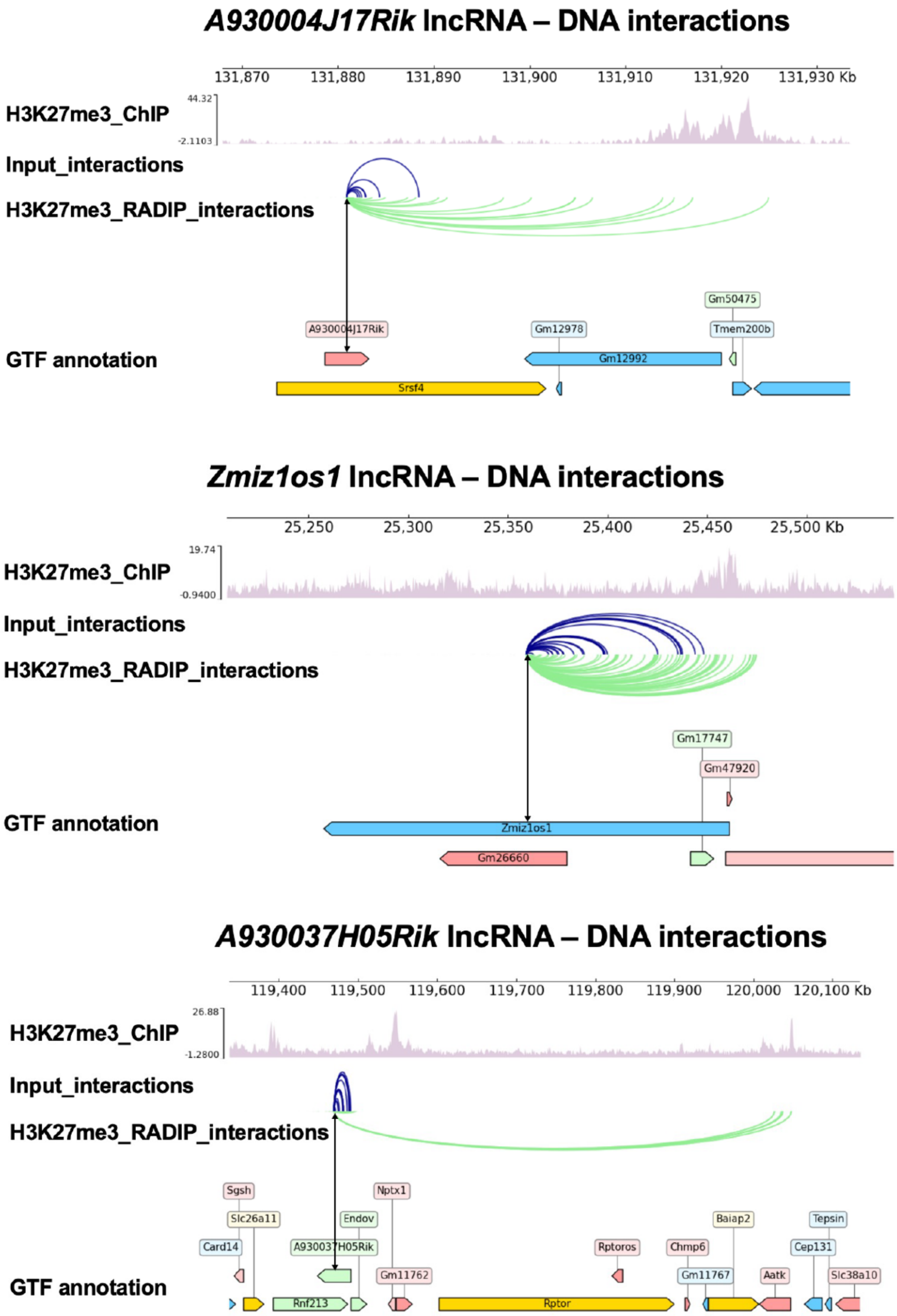


**Figure S4. H3K27me3-RADIP samples are enriched with lncRNAs interacting with H3K27me3 ChIP-seq peak DNA regions.**

*A930004J17Rik* lncRNA, *Zmiz1os1* lncRNA, and *A930037H05Rik* lncRNA were used as examples to show that the number of lncRNA–DNA interaction pairs was much greater in the H3K27me3-RADIP samples than in the Input samples. In addition, the target DNA tags of these lncRNAs in the H3K27me3-RADIP samples were in strong agreement with the H3K27me3 ChIP peaks.

Figure S5.

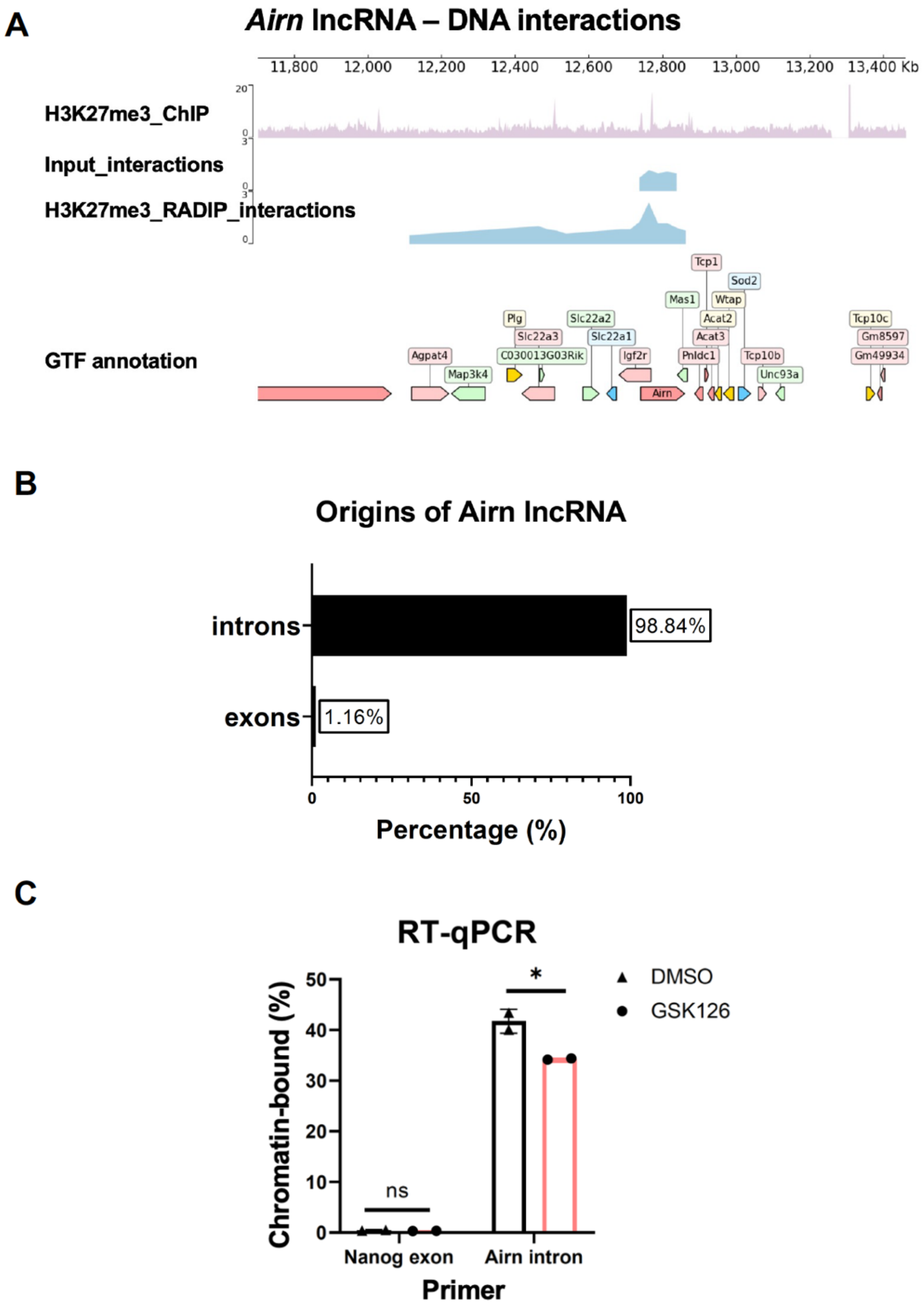


**Figure S5. The *Airn*–PRC2 complex is physically attached to chromatin.**

(**A**) H3K27me3-RADIP samples showed relatively high chromatin access of *Airn* lncRNA compared with Input samples, especially in the H3K27me3 ChIP-seq peak DNA regions. (**B**) Introns of *Airn* lncRNA were involved in almost all interactions with DNA. (**C**) The percentage of *Airn* lncRNA introns in the chromatin fraction was significantly reduced (**P*-value < 0.05) after treatment with GSK126, an EZH2 inhibitor. However, the percentage of *Nanog* mRNA exons used as a negative control did not change, indicating that some RNAs captured by H3K27me3-RADIP are physically attached to chromatin dependently of either the catalytic activity of EZH2 or H3K27me3 histone modification marks, although we do not know whether these RNAs are intron fragments, pre-mature mRNAs, or nascent RNAs.

### Figure S6.

**A**

chr.6

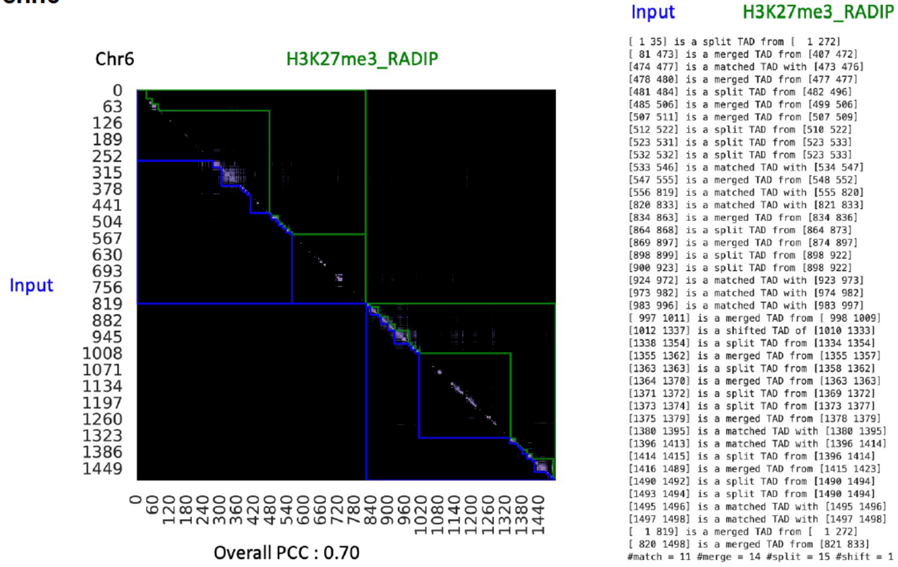


**B**

**RNAs related to common TLDs on chromosome 6 in H3K27me3_RADIP dataset (41 RNAs)**

Zfp777
Bola3
Dera
Tgfa
Mturn
Uba3
Pde3a
Sinhcaf
Bicd1
Hoxa11
Gm36640
Mthfd2
Slc4a5
Arl6ip5
Rbsn
Plekha8
Dennd5b
Zfp746
Gm28308
Pdia4
Frmd4b
Adcyap1r1
Rerg
Gcfc2
Tet3
Hoxaas3
Mob1a
Snrpg
Emp1
Resf1
Txnrd3
Gm765
Grin2b
Hoxa11os
Gm20696
Ptpro
Slc15a5
Ipo8
Cntnap2
Creb5
Hoxa10

**RNAs related to changed TLDs on chromosome 6 in H3K27me3_RADIP dataset (28 RNAs)**

Pde6h
Foxp1
Vps50
Uba3
Pde3a
Wipf3
Plekha8
Eogt
Gmcl1
Dennd5b
Jazf1
Sspo
Gm43948
Anxa4
Adcyap1r1
Rerg
Prickle2
Zfp282
Krba1
Aak1
Gm20696
Asns
Pdzrn3
Scrn1
Tmf1
Amn1
Eps8
Gm43950

**Figure S6. TLDs (topologically associating domain-like domains) for comparison of Input samples and H3K27me3-RADIP samples.**

(**A**) TLDs on chromosome 6 and the details of changes in TLDs on chromosome 6 for both types of samples are shown. (**B**) We identified and collected the anchor information for common TLDs and changed TLDs on chromosome (chr.) 6. We then determined which RNA species were related to these common TLDs (left) and changed TLDs (right) on chr. 6 in H3K27me3-RADIP samples as an example. Next, we followed this strategy for all chromosomes to determine the “Changed TLD” RNAs on a genome-wide level.

#### Figure S7.

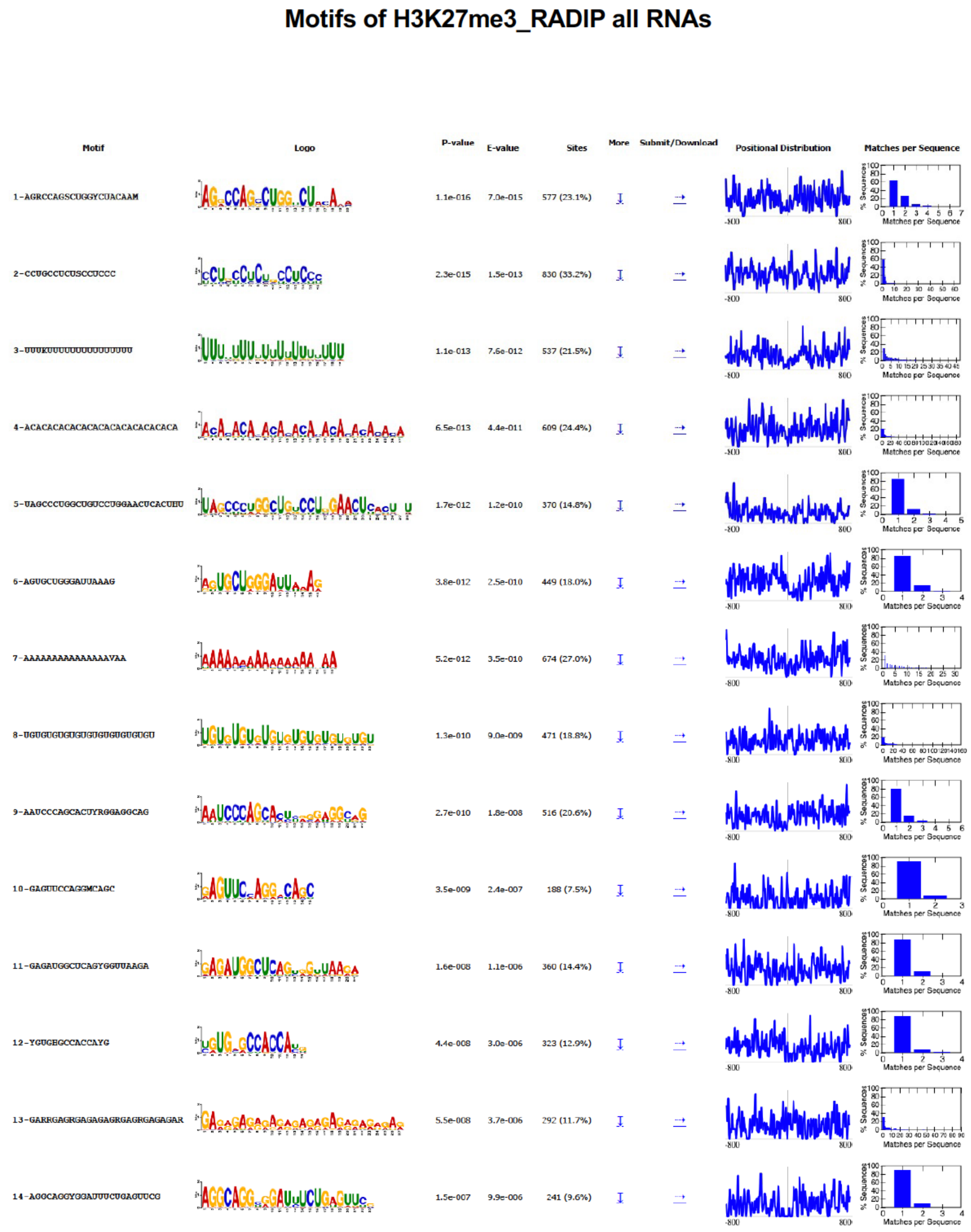


**Figure S7. Full list of enriched motifs from all RNA tags of H3K27me3-RADIP samples (page 1).**

### Figure S8.

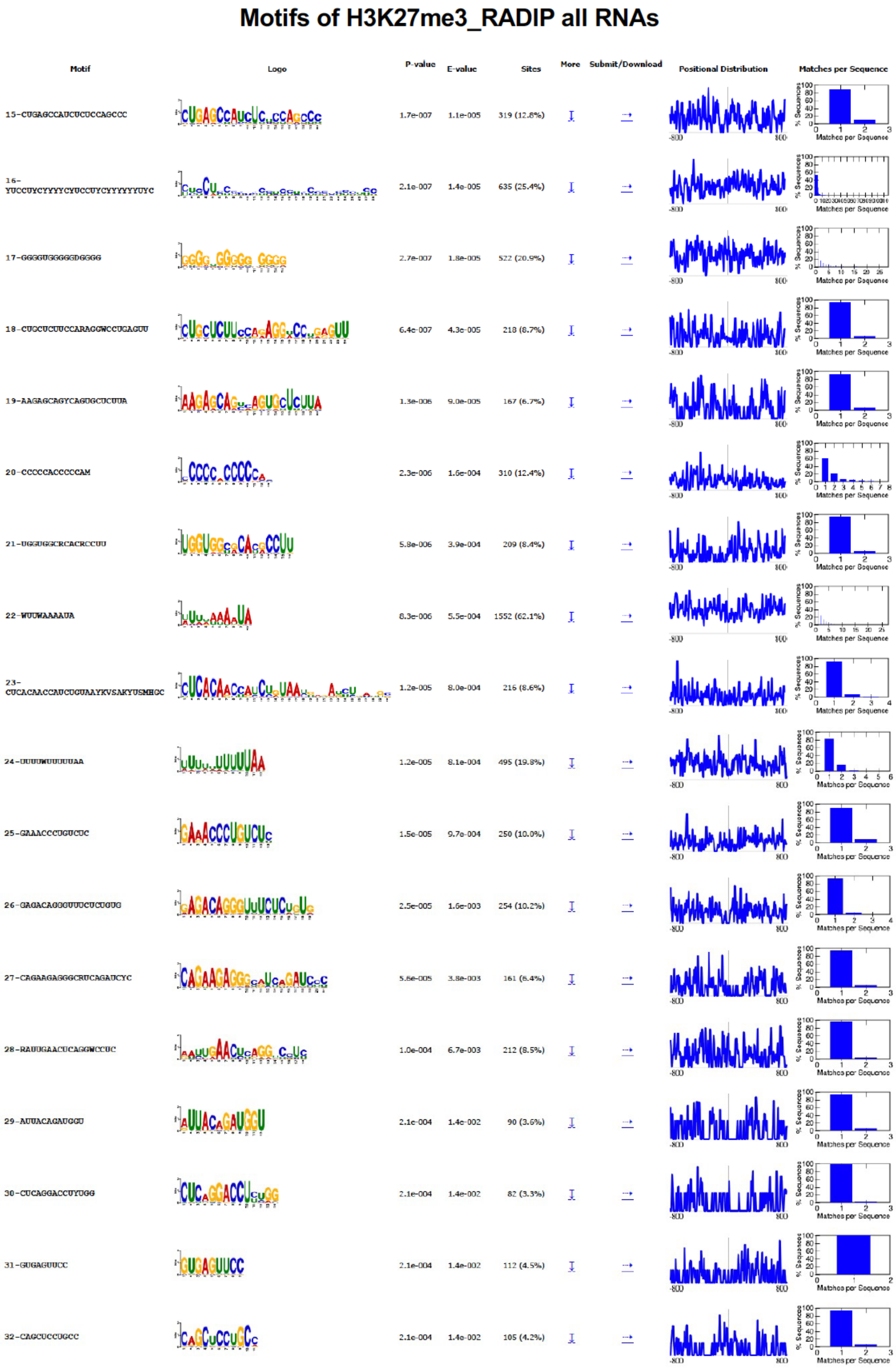


**Figure S8. Full list of enriched motifs from all RNA tags of H3K27me3-RADIP samples (page 2).**

### Figure S9.

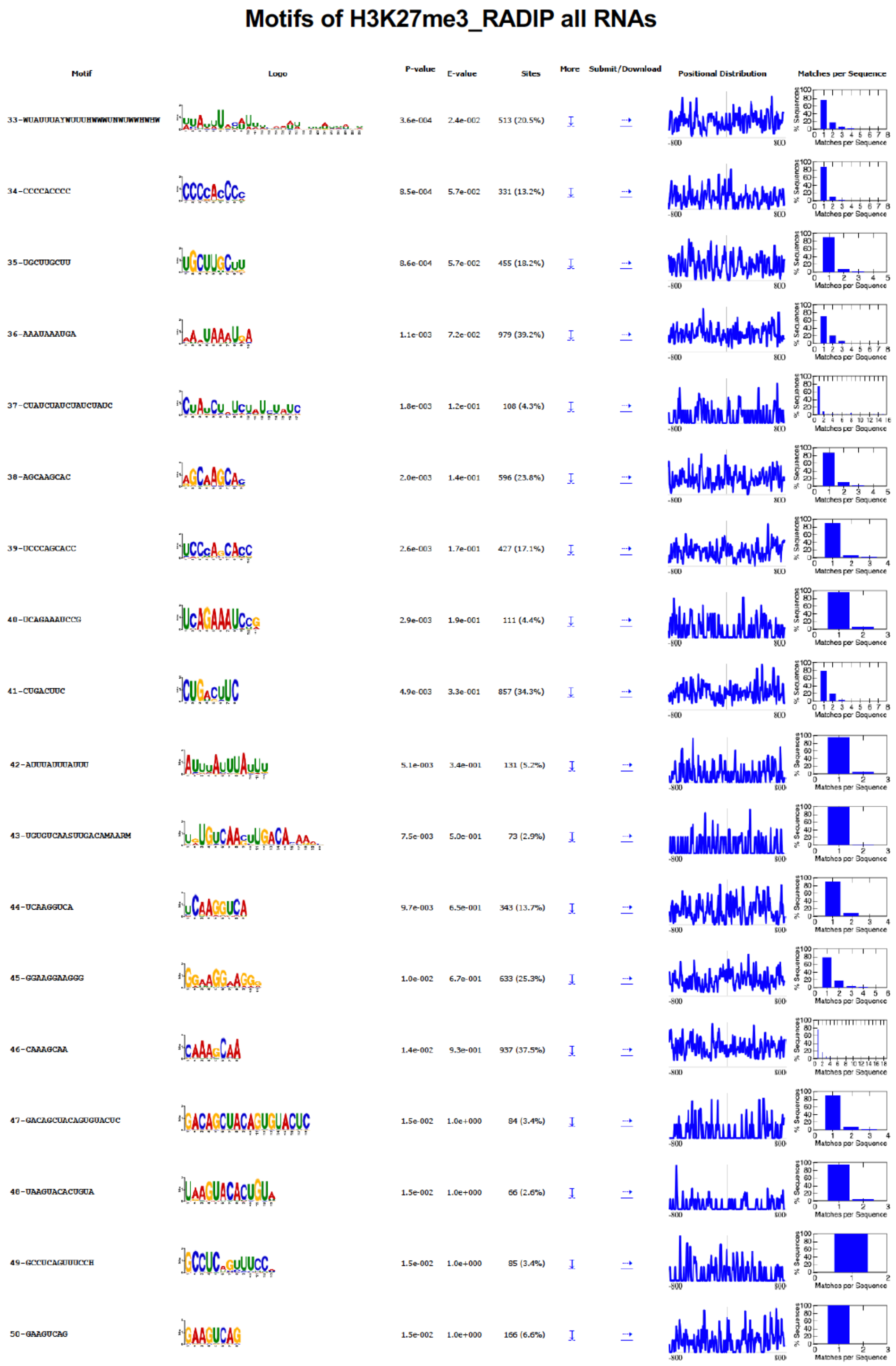


**Figure S9. Full list of enriched motifs from all RNA tags of H3K27me3-RADIP samples (page 3).**

#### Figure S10.

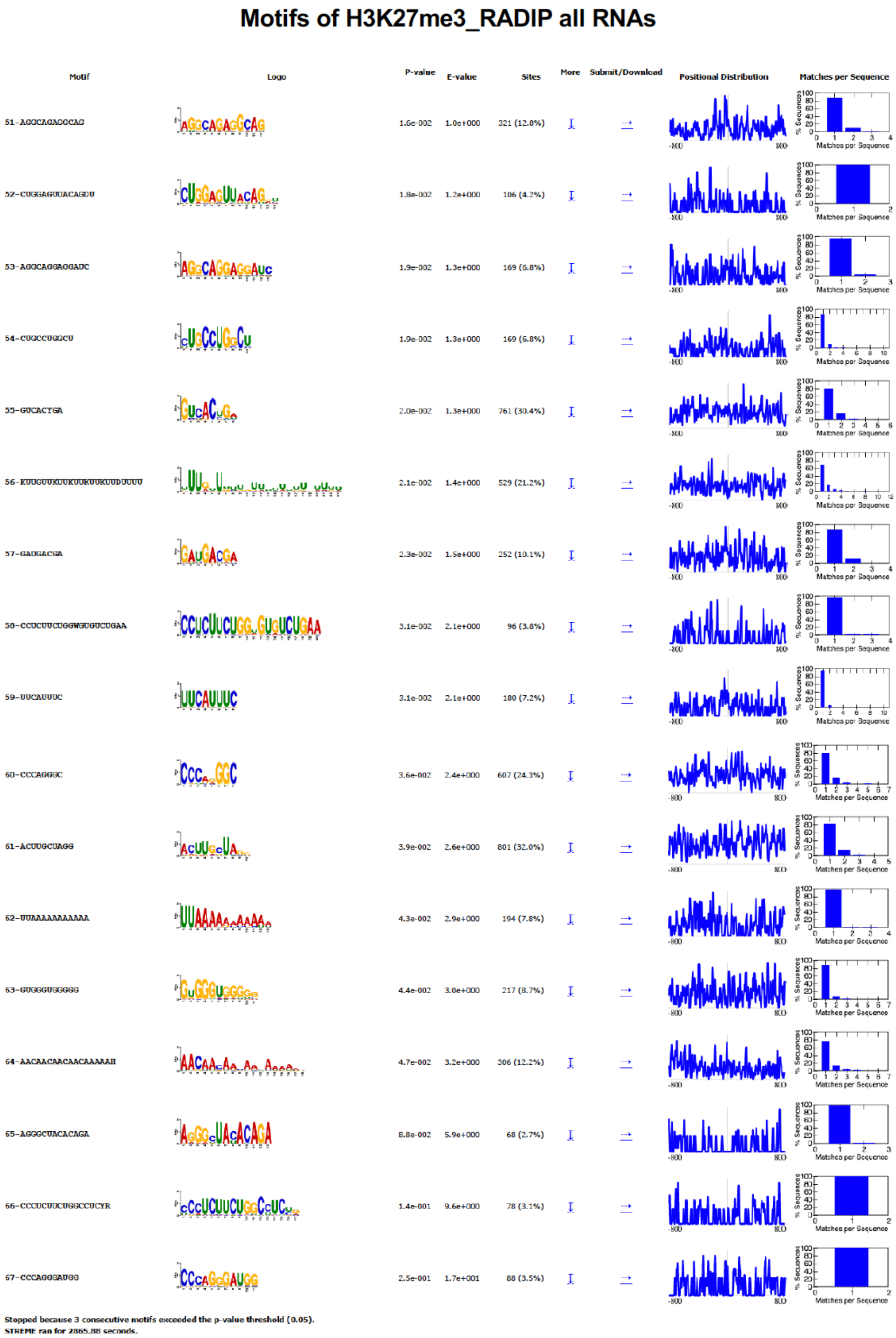


**Figure S10. Full list of enriched motifs from all RNA tags of H3K27me3-RADIP samples (page 4).**

**Figure S11.**

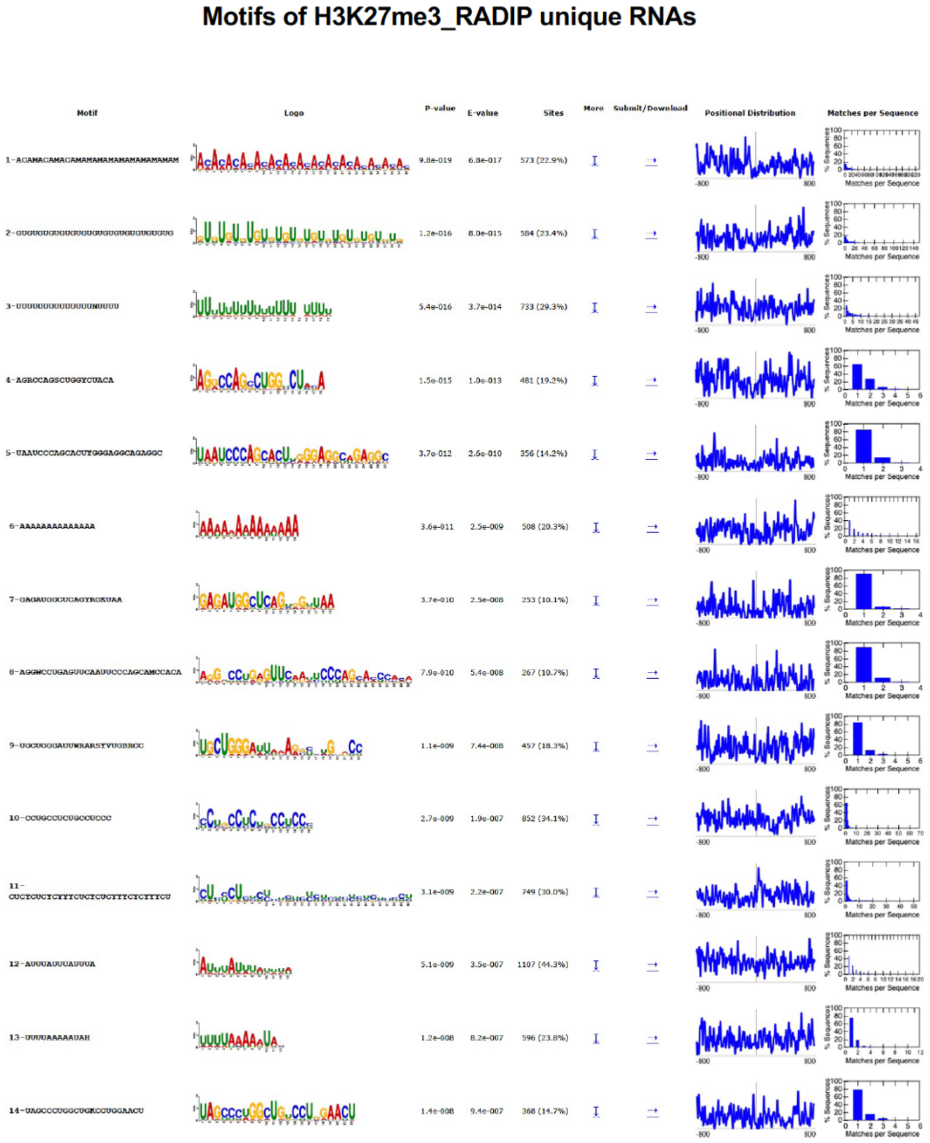


**Figure S11. Full list of enriched motifs from RNA tags unique to H3K27me3-RADIP samples compared with Input samples (page 1).**

#### Figure S12.

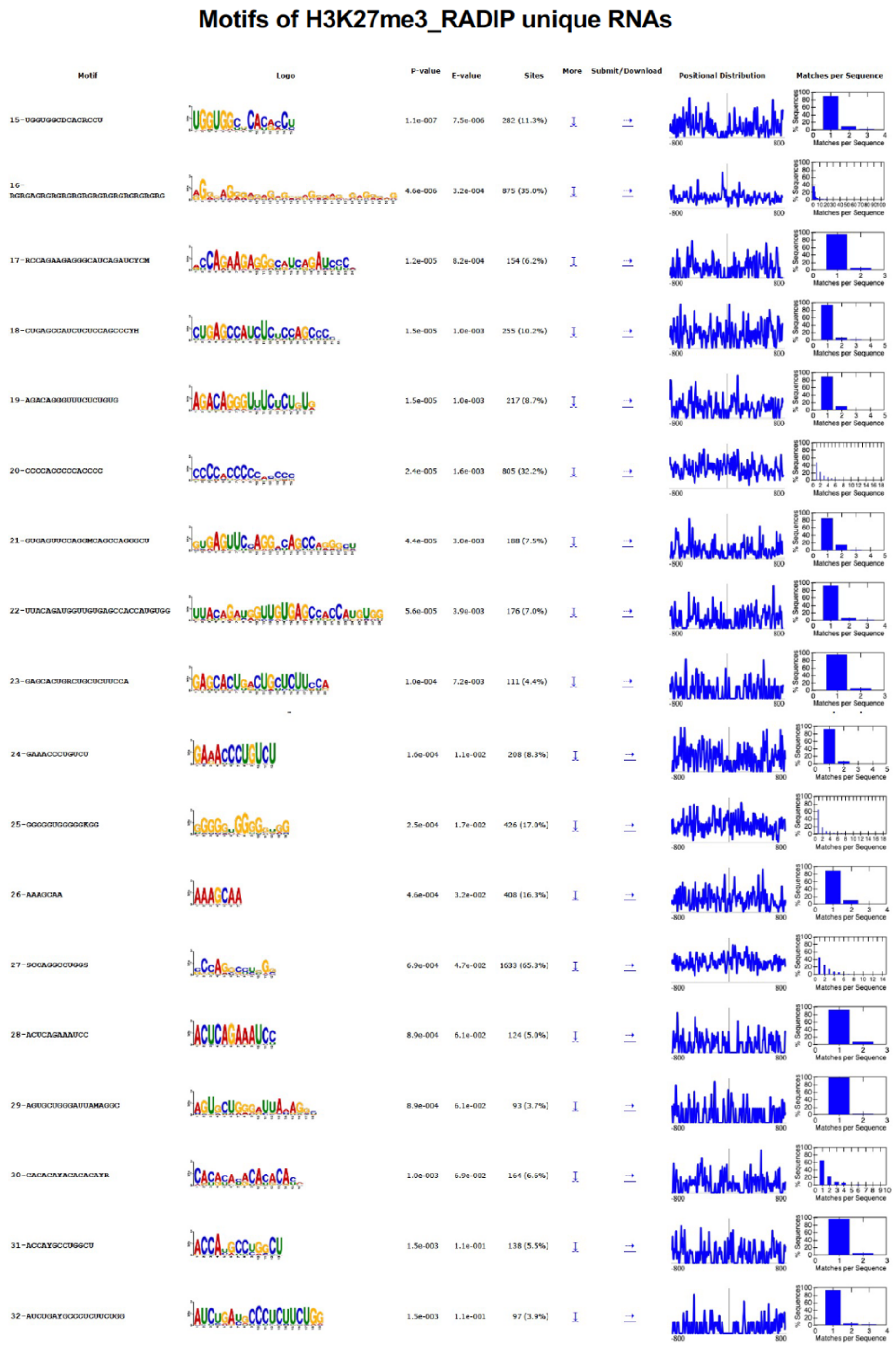


**Figure S12. Full list of enriched motifs from RNA tags unique to H3K27me3-RADIP samples compared with Input samples (page 2).**

#### Figure S13.

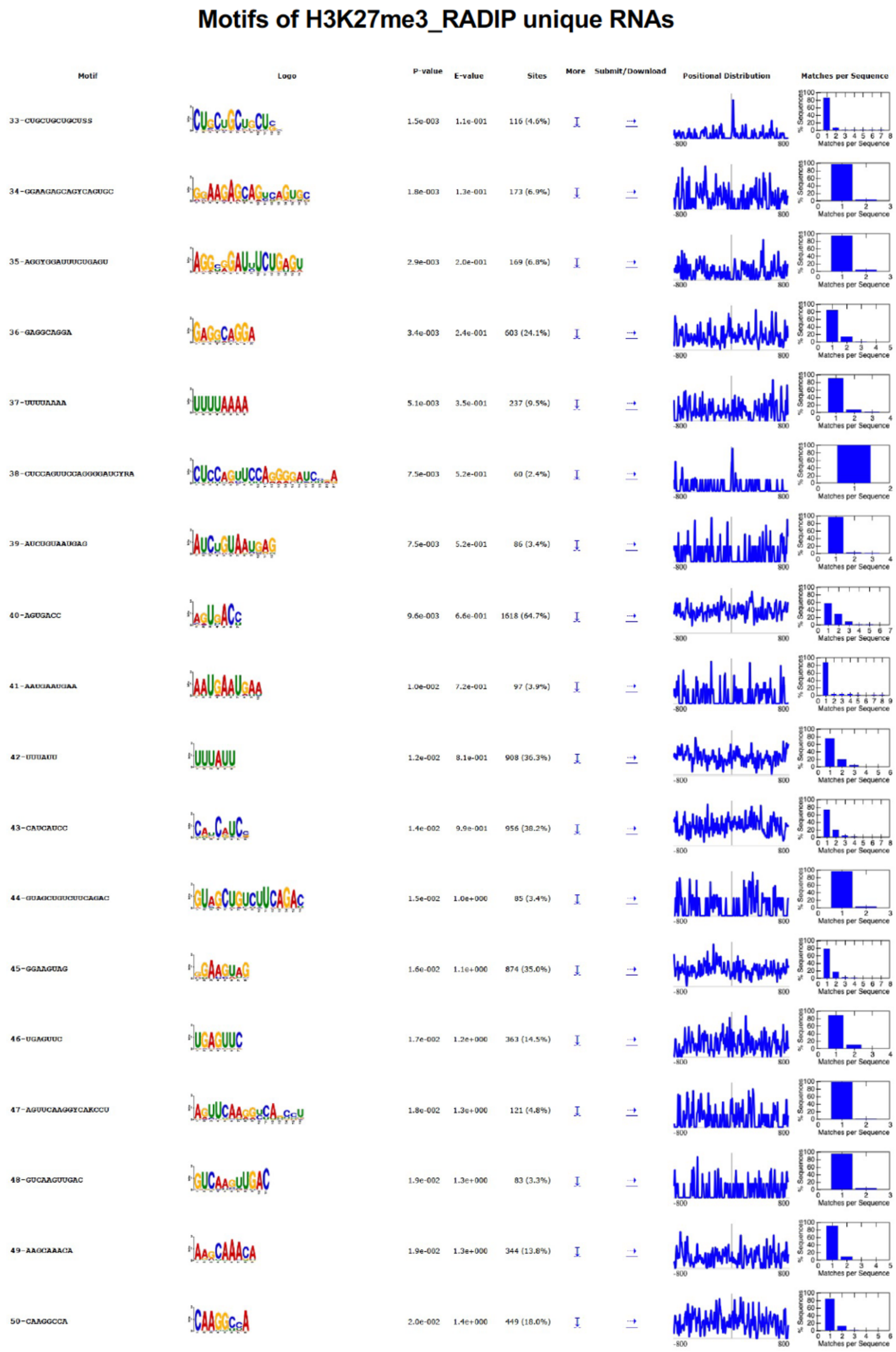


**Figure S13. Full list of enriched motifs from RNA tags unique to H3K27me3-RADIP samples compared with Input samples (page 3).**

#### Figure S14.

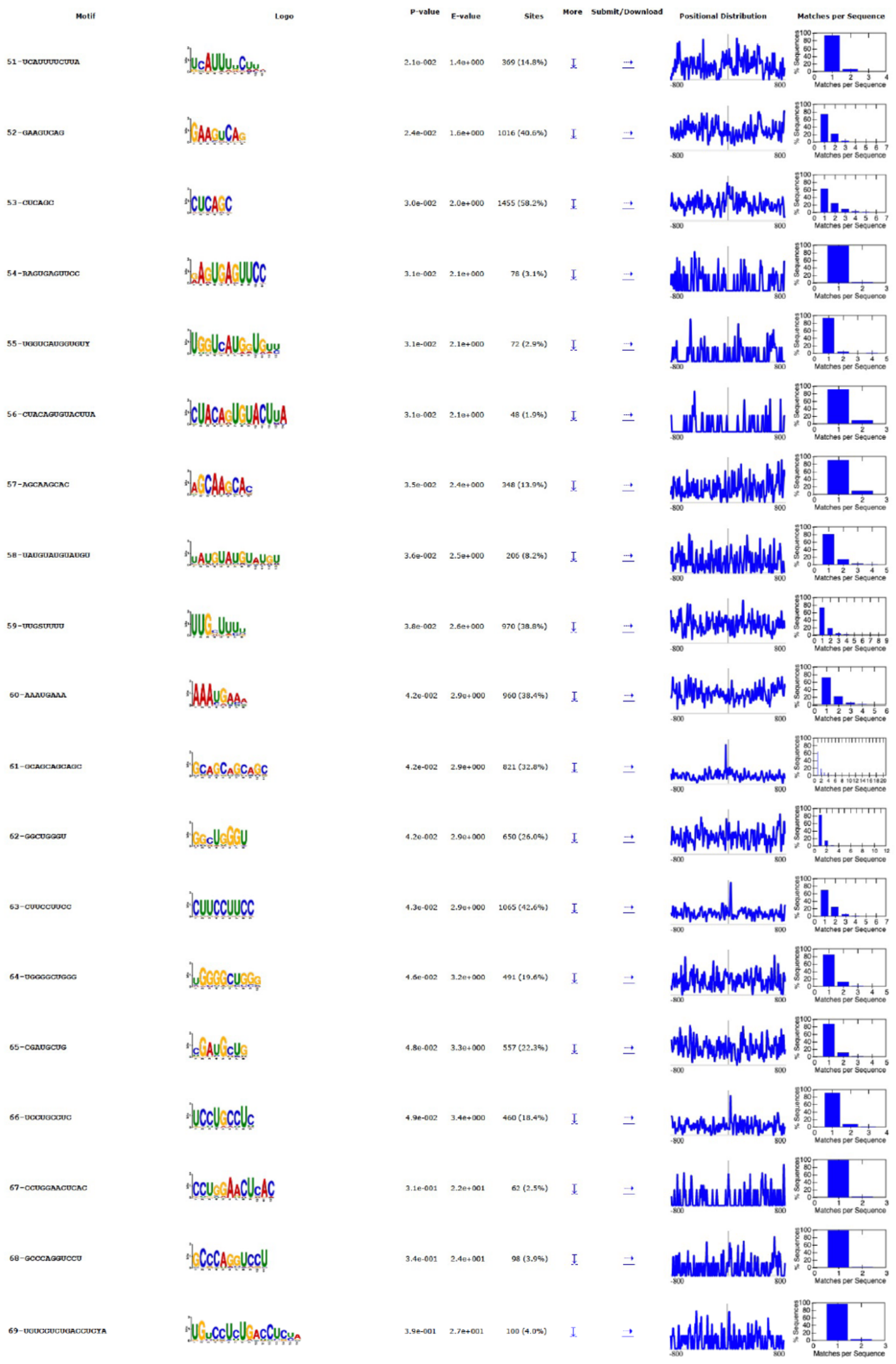

**Motifs of H3K27me3_RADIP unique RNAs**

| Motif | P-value | E-value | Sites |
|---|---|---|---|
| 51-UCAUUUUCUUA | 2.1e-002 | 1.4e+000 | 369 (14.8%) |
| 52-GAAGUCAG | 2.4e-002 | 1.6e+000 | 1016 (40.6%) |
| 53-CUCAGC | 3.0e-002 | 2.0e+000 | 1455 (58.2%) |
| 54-RAGUGAGUUCC | 3.1e-002 | 2.1e+000 | 78 (3.1%) |
| 55-UGGUCAUGGUGUY | 3.1e-002 | 2.1e+000 | 72 (2.9%) |
| 56-CUACAGUGUACUUA | 3.1e-002 | 2.1e+000 | 48 (1.9%) |
| 57-AGCAAGCAC | 3.5e-002 | 2.4e+000 | 348 (13.9%) |
| 58-UAUGUAUGUAUGU | 3.6e-002 | 2.5e+000 | 206 (8.2%) |
| 59-UUGSUUUU | 3.8e-002 | 2.6e+000 | 970 (38.8%) |
| 60-AAAUGAAA | 4.2e-002 | 2.9e+000 | 960 (38.4%) |
| 61-GCAGCAGCAGC | 4.2e-002 | 2.9e+000 | 821 (32.8%) |
| 62-GGCUGGGU | 4.2e-002 | 2.9e+000 | 650 (26.0%) |
| 63-CUUCCUUCC | 4.3e-002 | 2.9e+000 | 1065 (42.6%) |
| 64-UGGGGCUGGG | 4.6e-002 | 3.2e+000 | 491 (19.6%) |
| 65-CGAUGCUG | 4.8e-002 | 3.3e+000 | 557 (22.3%) |
| 66-UCCUGCCUC | 4.9e-002 | 3.4e+000 | 460 (18.4%) |
| 67-CCUGGAACUCAC | 3.1e-001 | 2.2e+001 | 62 (2.5%) |
| 68-GCCCAGGUCCU | 3.4e-001 | 2.4e+001 | 98 (3.9%) |
| 69-UGUCCUCUGACCUCYA | 3.9e-001 | 2.7e+001 | 100 (4.0%) |

Stopped because 3 consecutive motifs exceeded the p-value threshold (0.05).
STREME ran for 2801.29 seconds.

**Figure S14. Full list of enriched motifs from RNA tags unique to H3K27me3-RADIP samples compared with Input samples (page 4).**
