## Supplementary methods for "RADIP technology comprehensively identifies H3K27me3-mediated RNA-chromatin interactions"

*GSK126 treatment and cell fractionation.* Cells were treated with 0.1% DMSO (control) or 5 μM GSK126 (Merck) for 48 h. To extract the cytoplasmic, nuclear, and chromatin fractions, we followed a protocol described previously[1].

*RNA extraction, cDNA synthesis, and quantitative RT-PCR.* RNA was isolated using RNeasy Plus Mini Kit (Qiagen). An on-column DNase I (New England Biolabs) digestion was performed in accordance with the manufacturer's instructions. For chromatin RNA, we added 50 μL of precooled PBS to the chromatin pellet and resuspended the pellet by pipetting. We then added 500 μL of TRIzol™ Reagent (Invirtogen) with 5 μL of 0.5 M EDTA and incubated the mixture at 65 °C for about 10 min, with vortexing. Next, we added 100 μL of chloroform, vortexed the mixture briefly, and incubated it at room temperature for 5 min. This was followed by centrifugation for 15 min at 16,000*g* and 4 °C.

We transferred the upper aqueous layer to a new tube and added 3.5 sample volumes of Buffer RLT (Qiagen) We then followed the cleanup protocol of the RNeasy Kit. An equivalent volume of total RNA for each fraction was used for cDNA synthesis by using a SuperScript IV kit (Thermo Fisher Scientific) with Random 6-mer primers. For qRT-PCR, a PowerTrack SYBR Kit (Thermo Fisher Scientific) was used. The amount of RNA relative to that of HPRT (hypoxanthine phosphoribosyltransferase) in the cytoplasm fraction was calculated for each fraction, and then the percentage of RNA in the chromatin fraction was calculated.

The qRT-PCR primer sequences were as follows:
Airn_intron_Fw: GGGTCGAATCAGGTCACACA
Airn_intron_Rv: TCTCTGCTCTTGTGGGCAAA
Nanog_exon_Fw: TTCTGGGAACGCCTCATCAATGCCTGC
Nanog_exon_Rv: TCCTCAGGGCCCTTGTCAGCCTCAG
Hprt_exon_Fw: CAGGCCAGACTTTGTTGGAT
Hprt_exon_Rv: TTGCGCTCATCTTAGGCTT

*Details of modified CHiCANE method used to select significant pairs.* A modified negative binomial distribution approach, as implemented with the CHiCANE package in R, was used to calculate the adjusted *P*-value (*q* value) for each interaction pair. Then, only statistically significant interaction reads ($q$ value ≤ 0.1) were retained for subsequent analysis.

The purpose-defined CHiCANE-based method merely considers the intractability of the RNA tags. We used *trans* RNA–DNA interactions to produce the background.

Let $Y_{ij}$ denote the number of reads linking RNA tag $_i$ and target DNA tag $_j$, and let $t_i$ denote the number of reads linking the RNA tag $_i$ with all target DNA tags in *trans*. $\mu$ is a mean parameter, and $\theta$ is a dispersion parameter calculated from the count of all interaction pairs. After $\beta_0$, $\beta_1$, and the overdispersion term are fitted by the maximum likelihood method over the count of all interaction pairs, the MLEs (maximum likelihood estimates) of the coefficients $\beta_0$ and $\beta_1$ are used to provide an estimate of the new expected mean parameter $\mu_{ij}$ for each interaction pair, where:

$$\mu_{ij} = \beta_0 + \beta_1 \log(t_i + 1)$$

The MLE of the dispersion parameter $\theta$ is used as the dispersion parameter in the negative binomial model without zero observations (zero-count bins), where:

$$Y \sim \mathrm{NB}(\mu, \theta)$$

Therefore, $Y_{ij}$ is assumed to follow the fitted NB (negative binomial) regression with the adjusted mean parameter $\mu_{ij}$ and the adjusted dispersion parameter $\theta_{ij}$ for each interaction pair:

$$\mathrm{Var}(Y_{ij}) \sim \mu_{ij} + (\mu_{ij}^2 / \theta_{ij})$$

Next, a $P$ value for each interaction pair can be estimated for the observed counts $y_{ij}$ exceeding what is expected from the fitted NB model (background/noise), as:

$$P = P(Y_{ij} \geq y_{ij})$$

Finally, a multiple testing correction (by the Benjamini and Hochberg method) was applied to convert the $P$ value into a $q$ value (adjusted $P$ value or false discovery rate, or both) and identify significant interaction pairs that interacted more frequently than expected by chance, with a cutoff $q$ value of no more than 0.1. Only the significant datasets were used for subsequent analysis.

1. Sharma, H. et al. Decryption of sequence, structure, and functional features of SINE repeat elements in SINEUP non-coding RNA-mediated post-transcriptional gene regulation. *Nat Commun* **15**, 1400 (2024).
